# A Load-Induced Energetic Tipping Point Explains Selective Vulnerability of Substantia Nigra Neurons

**DOI:** 10.1101/2025.11.16.688654

**Authors:** Max Anfilofyev

## Abstract

Dopaminergic neurons in the substantia nigra pars compacta (SNc) are selectively vulnerable in Parkinson’s disease, whereas closely related neurons in the ventral tegmental area (VTA) are largely preserved. Many molecular stressors have been implicated—mitochondrial dysfunction, calcium load, -synuclein toxicity—but none by itself explains why two anatomically adjacent dopaminergic populations with similar molecular machinery diverge so sharply in their long-term fate. Here we analyze a **minimal two-variable energetic model** that captures only mitochondrial functional capacity, energetic reserve, and the combined structural and physiological loads imposed by axonal arborization and Ca^2^ handling.

The model reveals a **load-induced saddle-node bifurcation**: above a critical combined load, the system develops **coexisting high-energy and low-energy states**, separated by a narrow separatrix. VTA-like neurons (low arborization, modest Ca^2^ load) reside in a **monostable high-energy regime**, whereas SNc-like neurons (extreme arborization, elevated Ca^2^ demand) fall **inside a bistable window** under the representative calibration used here. In this regime, modest energetic insults can drive trajectories across the separatrix into an irreversible collapsed state, providing a compact dynamical explanation for long periods of stability followed by abrupt decline. We emphasize that this is a sufficiency result within a provisional calibration, not a quantitative claim that the exact experimental placement of every SNc and VTA neuron has already been validated.

Across parameter sweeps, two-parameter stability maps, and joint perturbation analyses, the bistable structure persists within the sampled nondimensional ranges. These results suggest that selective SNc vulnerability can arise from the **geometry of load-dependent energy–mitochondria coupling**, rather than from unique molecular defects, within this minimal framework and pending quantitative validation against bioenergetic measurements. The present model is spatially homogeneous and does not yet test whether the tipping point survives an explicit compartmental representation of the arbor; interventions shifting neurons away from this load-induced tipping regime may provide robust therapeutic leverage.

## 1. Introduction

Selective degeneration of substantia nigra pars compacta (SNc) dopaminergic neurons is a defining feature of Parkinson’s disease, yet dopaminergic neurons in the nearby ventral tegmental area (VTA) are comparatively resistant (15–17). This anatomical specificity remains difficult to reconcile with explanations centered on individual molecular stressors—mitochondrial impairment, oxidative stress, dopamine metabolism, or -synuclein aggregation—because these factors affect both populations and do not by themselves account for the stark difference in long-term outcomes (5–14,18–20).

A consistent observation across anatomical, physiological, and computational studies is that **SNc neurons carry extreme structural and metabolic loads**. Their axonal arbors span hundreds of thousands to millions of synapses (1–4,15–17,29,30), and prior energy-budget modeling has argued that propagating activity through such large dopaminergic arbors imposes a substantial ATP burden (31,32). Their pacemaking and calcium handling further couple electrical activity to mitochondrial demand and oxidant stress (5–9,33–37). These features suggest that SNc neurons operate closer to the limits of energetic feasibility than their VTA counterparts, raising the possibility that vulnerability reflects a **dynamical threshold** rather than a unique molecular defect.

Here we formalize this idea using a **minimal energetic model** with only two dynamical variables—energetic reserve *E* and mitochondrial functional capacity *M*—and two dimensionless load parameters representing axonal arborization *A* and Ca^2^ handling *C*. The model incorporates three empirically grounded interactions: mitochondria support energy; energy supports mitochondrial repair; and structural/functional loads increase both energetic consumption and mitochondrial vulnerability.

Despite its simplicity, this feedback loop is capable of generating a **load-dependent saddle-node bifurcation**. Below a critical load, the neuron has a single high-energy equilibrium. Above this threshold, the system contains two stable states—a healthy and a collapsed energetic state—separated by a saddle. The position of a neuron relative to this bistable window determines whether transient perturbations reliably recover or can trigger irreversible collapse.

Under the representative calibration used here, **VTA-like loads lie outside the bistable window**, implying recovery from perturbations in the model, whereas **SNc-like loads fall inside or near it**, positioning neurons near a separatrix where modest insults can drive abrupt decline. The resulting picture reframes selective vulnerability as a **geometric consequence of extreme anatomical and physiological load**, rather than requiring cell-type-specific molecular defects. We further show that the bistable region is robust across physiological parameter sweeps and two-dimensional parameter maps, indicating that the mechanism does not require fine-tuning within the explored parameter ranges.

The central goal of this work is therefore conceptual: to demonstrate that the **location of dopaminergic neurons in a load-induced energetic landscape** is sufficient to explain their divergent vulnerability patterns. This provides a unifying dynamical framework linking SNc anatomy, Ca^2^ physiology, mitochondrial strain, and the catastrophic collapse characteristic of Parkinsonian degeneration.

## 2. Minimal Energetic Model

### 2.1 Model Structure

To capture the essential energetic behavior of dopaminergic neurons without introducing unnecessary biochemical detail, we construct a **two-variable system** describing the interaction between:

- *E*(*t*): the cell’s *available energetic reserve*, normalized to [0, 1]. This variable integrates ATP availability, NADH balance, and the capacity of the cell to meet ongoing energetic demands (24,25).
- *M*(*t*): the cell’s *functional mitochondrial capacity*, also normalized to [0, 1]. This variable represents the collective ability of mitochondria to sustain oxidative phosphorylation under load, including turnover, repair, and stress-induced dysfunction (10–14,24).

SNc–VTA differences are introduced not by altering the equations themselves, but by adjusting two **load parameters** that modulate energy consumption and mitochondrial stress:

- *A*: the **axonal arborization load**, an effective load proportional to the energetic burden of maintaining and signaling through a large axonal arbor. Anatomically, SNc neurons have an arbor roughly 4–10× larger than VTA neurons, which we encode as higher values of *A* (1,3,15–17,29–32). The parameter *A* should not be read as synapse count itself; it is a nondimensional aggregate of synapse number, axon length, action-potential propagation cost, transmitter handling, and housekeeping burden.
- *C*: the **Ca**^**2**^ **-handling load**, representing the energetic overhead of pace-making, calcium buffering, and channel activity. In many simulations we set *C* = 1, with relative differences absorbed into the effective scaling of *A* (5–9,33–37). This coarse variable is intended to summarize total calcium-associated metabolic burden rather than any single conductance class. The combined load is modeled multiplicatively as *AC* for parsimony; this is a model-level assumption, not an independently validated law of dopaminergic physiology.

We treat *A* and *C* as time-independent on the timescales of the fast energetic dynamics analyzed here, reflecting the fact that axonal arborization and pace-making phenotype remodel over weeks to months rather than over the seconds-to-hours timescales on which *E* and *M* respond to metabolic insults. The model is therefore intended as a geometric description of a fast energy manifold embedded in slower structural context; if *A* or *C* drift slowly, singular-perturbation effects such as delayed tipping or rate-induced transitions could occur, but those effects are outside the present scope. The present manuscript does not analyze biological time series; it is a sufficiency result that isolates a minimal mechanism and leaves quantitative model–experiment validation to future work. A full prior-predictive or Bayesian calibration of the load parameters (*A, C*) and nonlinear gain *k*_2_ against independent data would be required to answer how often the SNc/VTA separation emerges without tuning the regime assignment to the conclusion.

The model structure (Figure 1) consists of three core interactions:

**Figure 1.**
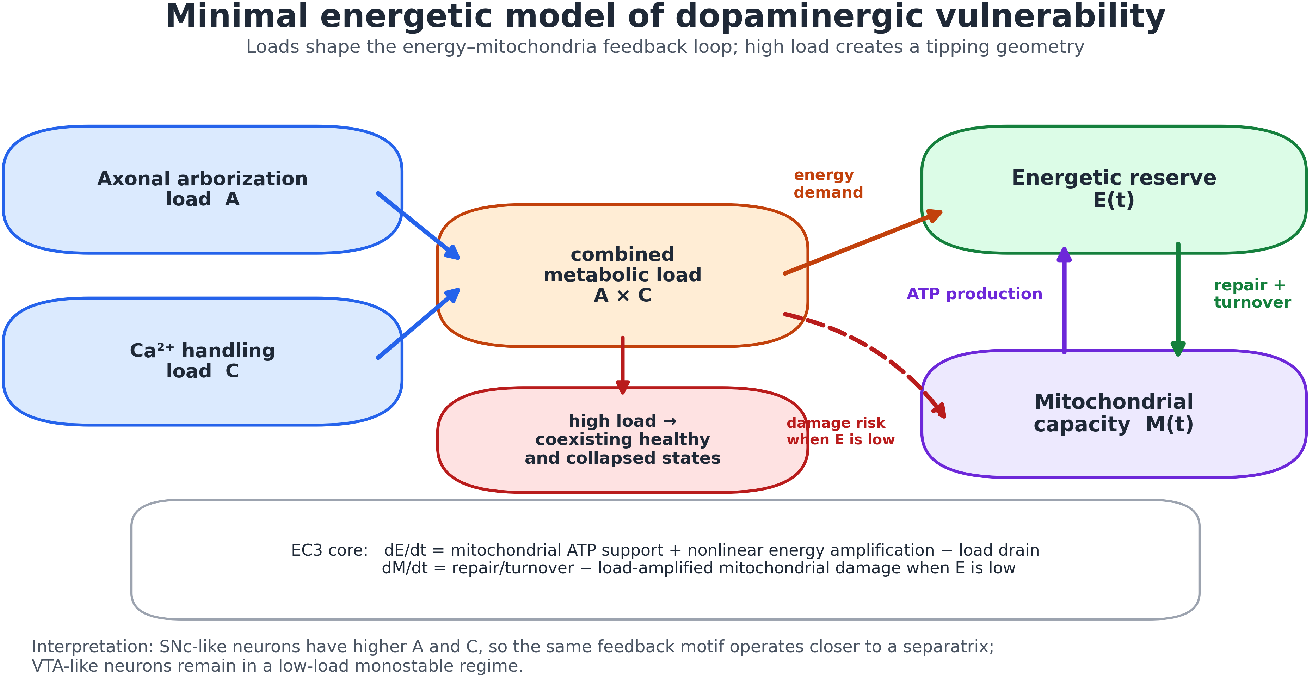
Minimal energetic model of dopaminergic neurons. Axonal arborization load *A* and Ca^2^ pacemaking load *C* combine into an effective metabolic demand that drains energetic reserve *E*. Mitochondrial capacity *M* supports ATP production and therefore increases *E*, while sufficient energy is required to maintain and repair mitochondria. When energetic reserve is low under high structural load, mitochondrial damage accumulates. This compact feedback loop—mitochondria support energy, energy maintains mitochondria, and loads destabilize both—is sufficient to generate coexisting healthy and collapsed energetic states under high load.

1. **Mitochondrial support of energy**. Functional mitochondria increase the energy reserve, reflecting ATP production proportional to available mitochondrial capacity (10–14,24).
2. **Load-dependent energy consumption**. Axonal and Ca^2^ loads drain energy at a rate that scales with both arbor size and Ca^2^-related metabolic demand (1,3,5–9,15).
3. **Energy-dependent mitochondrial maintenance**. Mitochondrial capacity is replenished through repair and turnover but is damaged when energetic reserve is insufficient to buffer load-induced stress, consistent with experimental observations in SNc neurons (10–14,15,24).

Together these interactions form a compact feedback loop: mitochondria support energy; energy enables mitochondrial maintenance; loads push both toward failure. Remarkably, this minimal structure is mathematically sufficient to generate **two coexisting energetic states**—a healthy, high-energy attractor and a collapsed, low-energy attractor—under high load. SNc neurons reside near the fold separating these states, whereas VTA neurons do not.

The vector field points inward on all boundaries of the normalized state domain [0, 1]^2^: at *E* = 0 energy production is nonnegative and points inward except at the (*E, M*) = (0, 0) corner; at *E* = 1 net flux is negative; at *M* = 0 mitochondrial repair is strictly positive; and at *M* = 1 turnover is nonpositive over the parameter ranges considered. Thus, all trajectories that begin in [0, 1]^2^ remain in this physiologically interpretable domain for all parameter values considered.

This parsimonious setup captures the qualitative behavior of dopaminergic neurons while keeping the model analytically transparent and numerically tractable within a normalized [0, 1]^2^ state space.

### 2.2 Equations and Qualitative Behavior

The interaction between energetic reserve *E* and mitochondrial capacity *M* produces a **nonlinear energy landscape** whose qualitative structure depends critically on axonal arborization load *A*. At low load, the system exhibits a single high-energy equilibrium. As load increases, the energy nullcline deforms until it develops two turning points—a geometry that permits three intersections with the mitochondrial nullcline and therefore the coexistence of two stable states and a saddle (21–23).

A natural concern is whether the nonlinear energetic amplification term *k*_2_*E*^2^(1 − *E*) is merely a mathematical convenience or reflects a biologically plausible mechanism. We introduced this term to capture the cooperative, saturating character of ATP-regenerating pathways documented at the cellular level. Oxidative phosphorylation exhibits nonlinear dependence on ADP and substrate availability, with sharp increases in ATP generation once mitochondrial membrane potential, NADH supply, and ADP levels cross critical thresholds, followed by saturation at high fluxes [24,25]. Similar input–output nonlinearities can arise from the combined action of dehydrogenases, substrate-channeling within mitochondrial microdomains, and feedback from ATP/ADP and AMP-activated protein kinase (AMPK) onto upstream glycolytic and tricarboxylic acid cycle steps [10–14,24,25]. Detailed mitochondrial ATP/Ca^2^ models often use mechanistic Michaelis–Menten, Hill-like, or coupled calcium-metabolism terms rather than this polynomial form (38–41). The specific polynomial form *E*^2^(1 − *E*) is therefore not intended as a detailed biophysical model of any single pathway, but as a minimal analytic surrogate that (i) is negligible at very low energetic reserve, (ii) increases superlinearly as *E* enters a cooperative regime, and (iii) saturates as *E* → 1. The present manuscript establishes a sufficiency result for this compact nonlinear feedback motif. In the current screening, simpler alternatives failed to reproduce the same fold geometry, so the result should be read as conditional on this surrogate choice rather than as a fully independent discovery from a broad unconstrained model class. Testing alternative saturating nonlinearities and fitting their parameters to dopaminergic-neuron bioenergetic data remain important robustness analyses for future work.

In the updated continuation analysis (Section 3), this fold structure emerges at a **left saddle-node** near *A* ≈ 0.86, beyond which a **bistable window** appears. Within this window—the interval

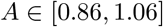

—the system supports:

1. a **high-energy attractor**,
2. a **low-energy attractor**, and
3. an **intervening saddle**, whose stable manifold forms the separatrix dirviding their basins.

Below *A* = 0.86, only the high-energy state exists; above *A* = 1.06, only the low-energy state remains.

Under this calibration, **VTA neurons** (arborization *A* ≈ 0.3–0.5) are **monostable** and robust to perturbations, whereas **typical SNc neurons** (*A* ≈ 0.9– 1.0) operate **inside the bistable regime**, precariously near the saddle’s stable manifold. SNc “outliers” with very large arbors (*A* ≈ 1.0–1.1) approach the **right fold**, where the high-energy and saddle equilibria annihilate, leaving collapse unavoidable under the model’s structure.

This establishes a clear dynamical principle: **axonal load determines whether the neuron lives in a monostable or bistable energetic regime**, with Ca^2^-handling load acting as a multiplicative co-load that shifts the same saddle-node geometry (Supplementary S3.3). In the main figures we therefore hold *C* = 1 to isolate the anatomical contribution of *A*, and treat *C* as a **multiplicative co-load** that effectively rescales *A*; their **combined** burden, *AC*, determines whether a neuron lies inside or outside the bistable window in parameter space. **A full** (*A, C*) **stability map (Supplementary Figure S8) confirms this picture: the bistable region appears as an oblique band in** (*A, C*) **space. Along the** *C* = 1 **slice used in the main text, VTA-like neurons at** (*A, C*) = (0.40, 1.0) **fall in the monostable region, whereas SNc-like neurons at** (*A, C*) = (1.00, 1.0) **lie inside the bistable band. Increasing Ca**^**2**^ **-handling load** *C* **shifts this band leftward (inducing bistability at lower** *A***), while decreasing** *C* **shifts it rightward, consistent with** *A* **and** *C* **acting as multiplicative co-loads**. The current mapping from morphology and physiology into (*A, C*) is therefore a provisional calibration for the SNc/VTA comparison, not an independently blinded fit.

To illustrate the qualitative structure of the energy dynamics, Figure 2A shows a conceptual landscape representation of the system’s two attractors and the saddle separating them. The high-energy attractor corresponds to normal physiological operation, while the low-energy attractor represents an irreversible collapsed state. The saddle acts as the basin boundary: perturbations that remain on the healthy side of this boundary recover, whereas perturbations that cross it transition to the collapsed state. This geometric picture anticipates the mathematical structure revealed in the phase-plane analysis (Figure 2B) and highlights why neurons experiencing large structural loads, such as substantia nigra dopaminergic neurons, reside precariously close to the tipping boundary, whereas ventral tegmental area neurons lie far from the saddle and therefore exhibit robust recovery after metabolic or calcium-related stress (15–17,21–23).

**Figure 2A.**
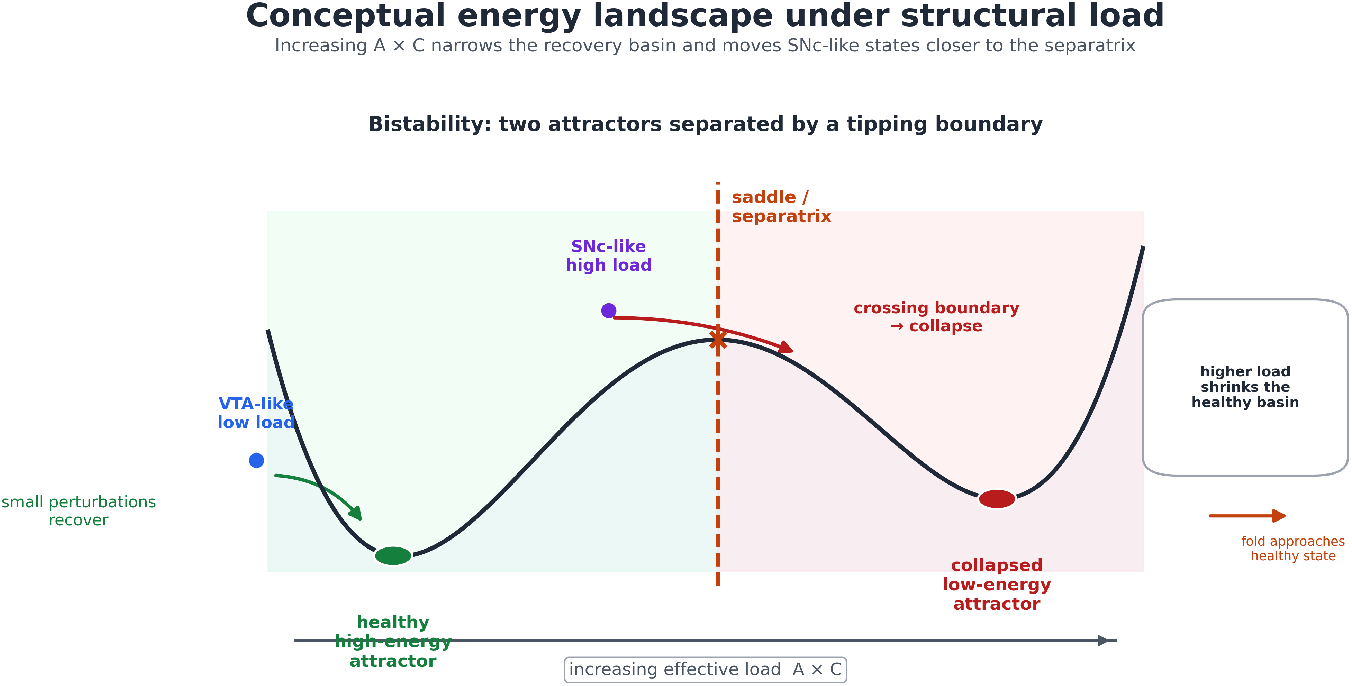
Conceptual energy landscape illustrating bistability under structural load. The minimal energetic model predicts two coexisting stable states separated by a saddle (tipping boundary). The upper basin corresponds to a healthy, high-energy attractor, while the lower basin represents an energetically collapsed state. Small perturbations within the healthy basin are restored, whereas sufficiently large perturbations cross the separatrix and drive the system into the collapsed attractor. Increasing axonal load *A* shifts the saddle leftward and shrinks the healthy basin. VTA neurons, which experience low load, lie deep within the healthy basin; SNc neurons, under high load, lie near the saddle and are therefore vulnerable to tipping into collapse (15–17,21–23). Under the updated bifurcation structure, VTA neurons (*A* ≈ 0.3–0.5) lie left of the saddle-node fold at *A* ≈ 0.86, whereas SNc neurons (*A* ≈ 0.9–1.0) lie inside the bistable window for which this landscape applies.

**Figure 2B.**
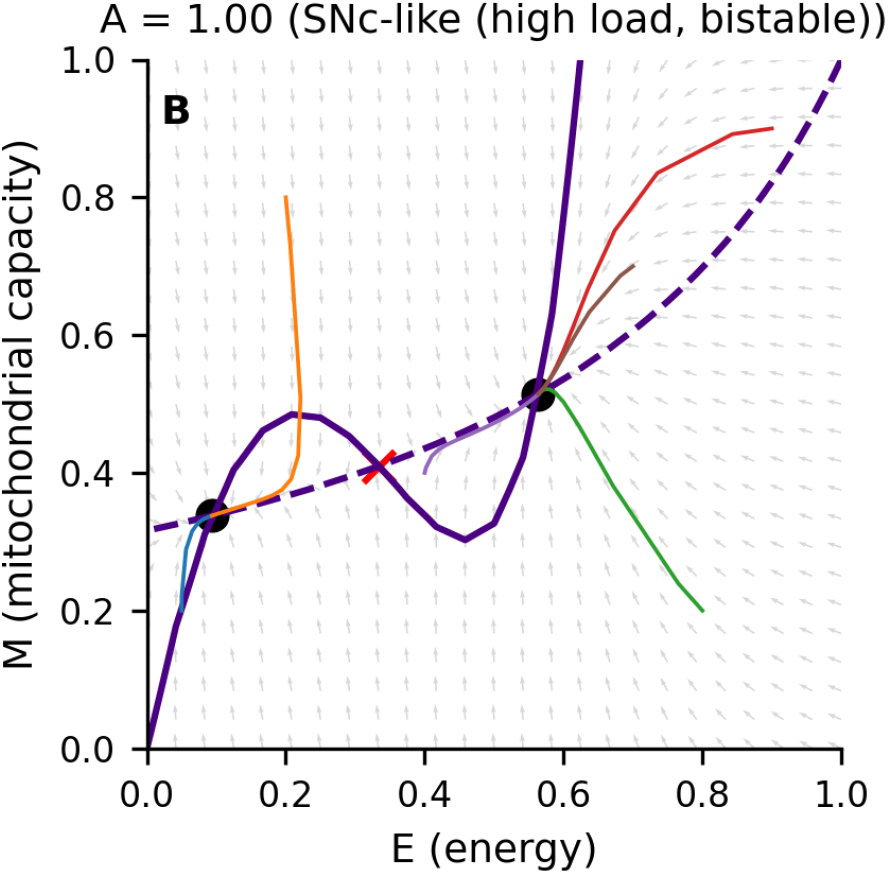
Phase portrait at a substantia nigra–like load (*A* = 1). Grey arrows show the vector field in the (*E, M*) plane. The solid purple curve denotes the energy nullcline (*dE*/*dt* = 0), forming an S-shaped curve with two folds, and the dashed purple curve denotes the mitochondrial nullcline (*dM*/*dt* = 0). Their intersections yield three equilibria: a high-energy stable fixed point (upper black dot), a low-energy stable fixed point (lower black dot), and an intermediate saddle (red cross). Colored trajectories illustrate that initial conditions on one side of the saddle’s stable manifold converge to the healthy high-energy attractor, whereas those on the other side collapse into the low-energy state. This phase-plane geometry provides a concrete dynamical realization of the conceptual landscape in Figure 2A and shows that SNc-like loads place the system inside a load-induced tipping regime (21–23).

This geometry is the mathematical signature of a **tipping point**: a minimal energetic failure mode that emerges directly from load-dependent energy consumption and energy-dependent mitochondrial stress (21–23). In subsequent sections, we show that ventral tegmental area neurons reside far from this fold, while substantia nigra neurons sit near it, making them uniquely susceptible to collapse (15–17).

## 3. Bifurcation Analysis: Load-Driven Emergence of a Tipping Point

To determine how axonal arborization load shapes the neuron’s energetic stability, we performed a one-parameter continuation in the load parameter *A* and computed all equilibria across the dopaminergic range. As shown in Figure 3, the system exhibits a **saddle-node bifurcation** in energetic reserve *E*, separating a low-load **monostable** regime from a higher-load **bistable** regime in which healthy- and collapsed-energy states coexist.

**Figure 3.**
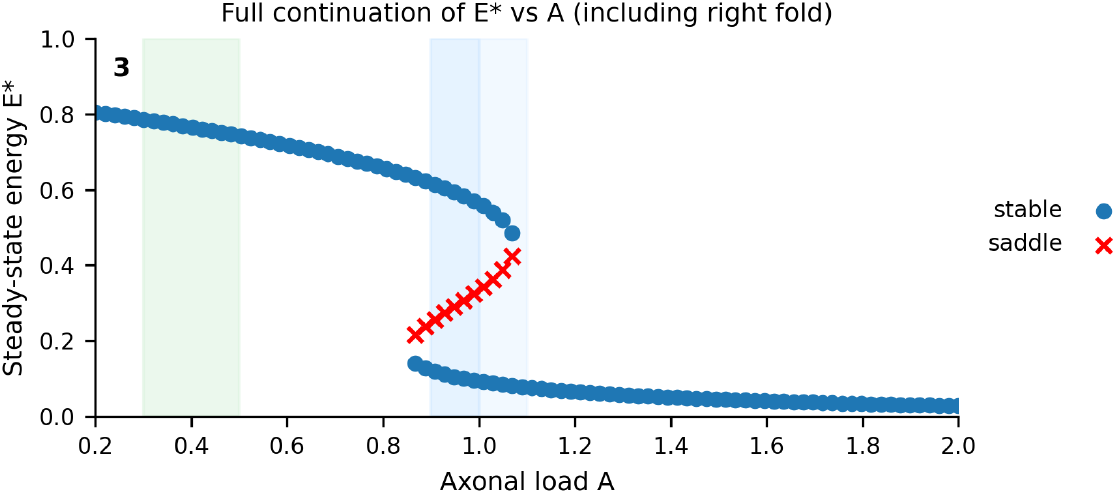
Saddle-node bifurcation of energetic reserve as a function of axonal load. The steady-state energetic reserve *E*^*^ is plotted as a function of axonal load *A*. Solid markers denote stable equilibria; crosses denote saddles. Vertical shaded bands indicate VTA-like loads (*A* ≈ 0.3–0.5), typical SNc loads (*A* ≈ 0.9–1.0), and the SNc high-outlier/uncertainty band (*A* ≈ 1.0–1.1).

**Figure 4.**
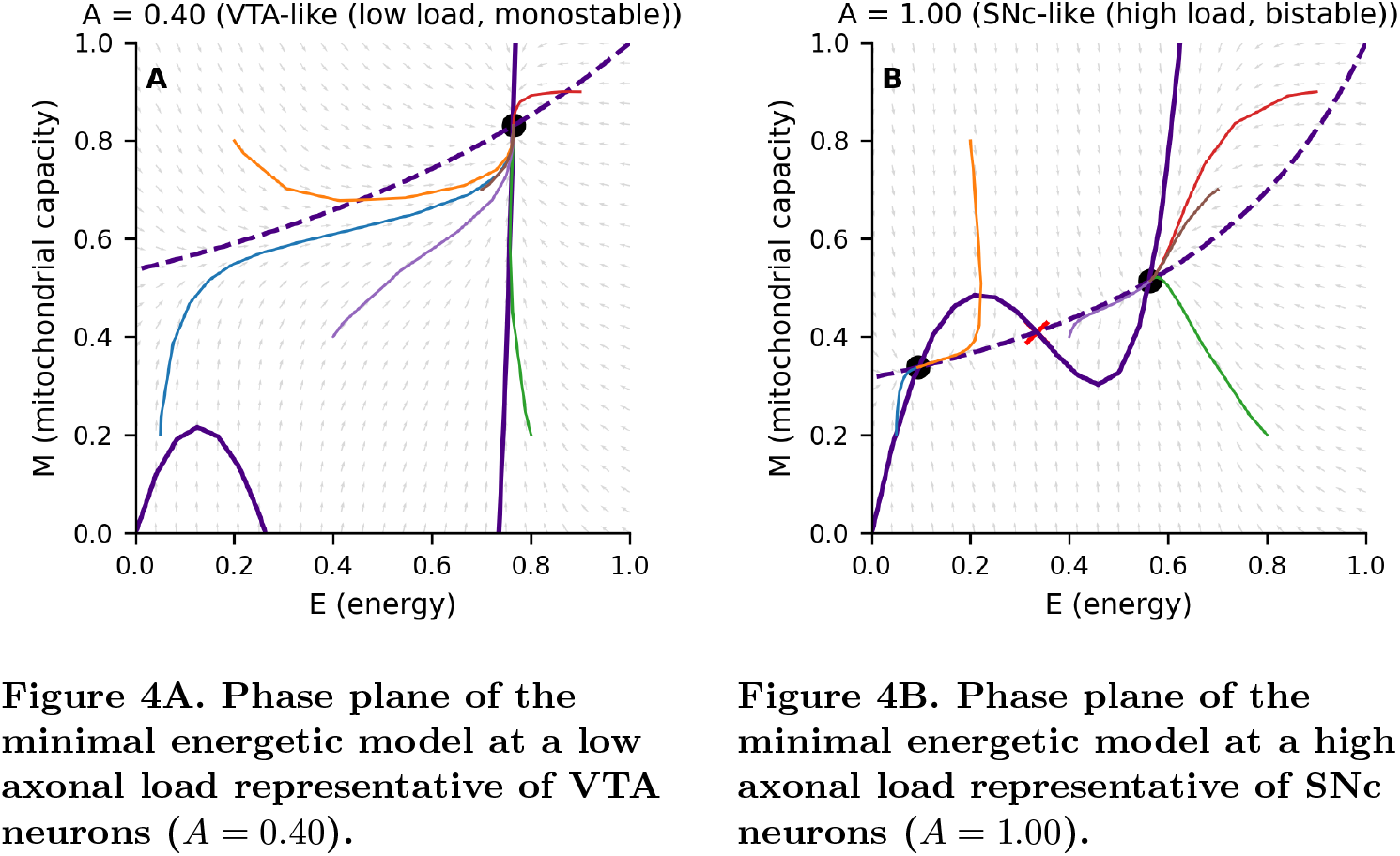
Phase-plane comparison of VTA-like and SNc-like dopaminergic neurons. *Left (Figure 4A):* The horizontal axis shows energetic reserve *E* and the vertical axis mitochondrial capacity *M*. Gray arrows denote the vector field, the solid curve the energy nullcline (*dE*/*dt* = 0), and the dashed curve the mitochondrial nullcline (*dM*/*dt* = 0). In this low-load regime the null-clines intersect only once, at a high-energy, high-mitochondrial-capacity equilibrium. Sample trajectories (colored curves) initiated from widely separated initial conditions—including states with low energy and/or impaired mitochondria— are all attracted to this single fixed point. The absence of additional fixed points or a separatrix indicates a globally attracting, monostable high-energy regime, in which VTA-like neurons robustly recover their energetic state following transient perturbations rather than tipping into collapse. Under the updated bifurcation structure, VTA neurons (*A* ≈ 0.3–0.5) lie left of the saddle-node fold at *A* ≈ 0.86, whereas SNc neurons (*A* ≈ 0.9–1.0) lie inside the bistable window for which this landscape applies. *Right (Figure 4B):* As in the left panel, the horizontal axis shows energetic reserve *E* and the vertical axis mitochondrial capacity *M*. The solid curve denotes the energy nullcline (*dE*/*dt* = 0) and has an S-shaped profile, while the dashed curve shows the mitochondrial nullcline (*dM*/*dt* = 0). At this high load the nullclines intersect three times: at a low-energy, low-mitochondrial-capacity fixed point, a high-energy fixed point, and an intermediate saddle (red marker). Sample trajectories (colored curves) initiated on either side of the saddle’s stable manifold diverge toward different long-term outcomes: those starting above and to the right of the separatrix relax to the high-energy attractor, whereas those starting below or to the left are drawn into the low-energy attractor. The coexistence of these two energetic fates under the same parameter set, separated only by a narrow dynamical boundary in this spatially homogeneous whole-cell model, illustrates how extreme structural load places SNc-like neurons inside a bistable regime where modest perturbations can tip them from normal function into irreversible energetic collapse (1,3,5–9,15–17,21–23).

At low axonal load—corresponding to the **ventral tegmental area (VTA)** range—the system contains a **single stable equilibrium**, a robust high-energy attractor. As *A* increases, the energy nullcline bends downward, and at a critical load value near

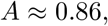

the system undergoes a **left saddle-node fold** that gives rise to a **low-energy stable state** and an intervening **saddle point** (21–23). This marks the onset of **bistability**. For loads

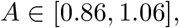

the system contains three equilibria: a healthy high-energy attractor, a collapsed low-energy attractor, and a saddle separating their basins.

Crucially, the **typical SNc load range (***A* ≈ 0.9**–**1.0**) lies entirely to the right of the left fold**, meaning that SNc neurons operate **inside** the bistable regime rather than approaching it from below. Even modest metabolic or calcium-driven perturbations can push the system across the saddle’s stable manifold and into the collapsed state, providing a compact explanation for the long periods of apparent resilience followed by sudden, irreversible decline in substantia nigra dopaminergic neurons (1,3,5–9,10–17,24).

At higher load the bistable structure eventually terminates: the high-energy and saddle branches coalesce in a **right fold** near

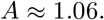

This value lies **above the typical SNc range** (*A* ≈ 0.9–1.0) but within the broader **SNc outlier/uncertainty band** (*A* ≈ 1.0–1.1). Within this minimal model, recovery from the low-energy state by changing load alone would require reducing *A* below the left fold (approximately 0.86) to re-enter the monostable high-energy regime; absent implausibly large changes in arborization or concurrent shifts in other parameters (e.g., *k*_*M*_, *L*_1_, or *C*), axonal pruning alone therefore appears unlikely to restore the healthy state once collapse has occurred.

Extended parameter sweeps and numerical continuation analyses (Supplementary Figures S1–S6) confirm that this saddle-node structure is a **robust feature** of the model under one-at-a-time parameter variations: the location of the folds shifts modestly with changes in mitochondrial turnover, Ca^2^-handling cost, and energy-dependent mitochondrial damage, but the bistable window persists (21–23). Because the nonlinear energetic amplification term *k*_2_ is empirically unconstrained, we also conducted a two-parameter sweep across *A* and *k*_2_ (Supplementary Figure S11). The bistable region persisted across a broad range of *k*_2_ values (4–9), with the diagonal geometry preserved. This indicates that bistability does not depend on fine-tuning nonlinear metabolic feedback, but is instead a structurally stable property of the model class.

To examine how axonal load *A* interacts with Ca^2^-handling load *C*, we computed a two-parameter stability map of the EC3 model (Supplementary Figure S8). The system exhibits a smooth, diagonally oriented bistable band in (*A, C*) space: lower *C* values shift the onset of the saddle-node to higher *A*, while higher *C* values shift it leftward. Importantly, VTA-like loads (*A* ≈ 0.4) remain monostable across the entire physiologically plausible range of *C*, whereas SNc-like loads (*A* ≈ 1.0) fall squarely inside the bistable band unless *C* is markedly reduced. Thus, Ca^2^ burden modulates but does not eliminate the load-driven energetic tipping point.

Although a load-induced saddle-node bifurcation is **sufficient** to generate abrupt and effectively irreversible collapse in this framework, we do not claim that it is the **unique** minimal mechanism capable of doing so. Other compact model classes—such as slowly drifting parameters or noise-driven transitions in monostable systems—can in principle yield superficially similar failure dynamics, and distinguishing among these alternatives will require targeted experiments.

Together, these results demonstrate that **axonal arborization and calcium-handling demand jointly govern dopaminergic energetic stability**. In the main text we use *A* as the explicit continuation parameter and set *C* = 1 to isolate anatomical differences. Supplementary sweeps and a full (*A, C*) stability map (Supplementary Figure S8) show that changes in *C* act as an effective rescaling of load: their **combined** burden *AC* determines whether a neuron lies inside or outside the bistable band. Along the *C* = 1 slice, the folds at *A* ≈ 0.86 and *A* ≈ 1.06 match those in Figure 3; at higher *C* the same folds appear at lower *A*, and at lower *C* they shift to higher *A*.

## 4. Comparison of Substantia Nigra and Ventral Tegmental Area Neurons

A defining characteristic of substantia nigra pars compacta (SNc) dopaminergic neurons is their extraordinarily large and widely distributed axonal arbor. Anatomical reconstructions indicate that a single SNc neuron forms hundreds of thousands to millions of synapses, a structural scale unmatched by most other neuronal types (1,3,15,16). In contrast, dopaminergic neurons in the ventral tegmental area (VTA) innervate far fewer targets, with substantially smaller arbor size and reduced calcium-handling burden during pacemaking (5–9,15– 17). These anatomical and physiological differences map naturally onto the load parameter *A* in our model, with *C* capturing additional Ca^2^ burden.

Updated bifurcation results place **VTA neurons** (*A* ≈ 0.3–0.5) **entirely left of the left fold** at *A* ≈ 0.86, meaning they live in a **strictly monostable** portion of parameter space. Their phase planes therefore contain **only a single high-energy attractor**, with no saddle and no collapsed state available. **Supplementary Figure S8 shows that this remains true across the full range of Ca**^**2**^ **loads considered (***C* ∈ [0.5, 1.5]**): for all** *C* **in this band, VTA-like points lie in the monostable region, whereas SNc-like points cluster within the bistable band**.

**SNc neurons**, by contrast, have typical arborization loads of

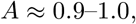

which lie **inside the bistable window** identified in Section 3. Their phase planes show:

- a high-energy attractor,
- a low-energy attractor,
- and the saddle separating the two basins.

SNc neurons therefore operate **on the interior of a bistable region**, not merely near its edge. This matters because perturbations need not be large to reach the saddle’s stable manifold.

A small subset of SNc neurons—those with unusually large arbors—likely fall in

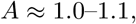

a band that approaches the **right fold** at *A* ≈ 1.06, where the healthy state disappears entirely. These “outliers” are predicted to be the **most vulnerable** neurologically, consistent with observed heterogeneity in SNc degeneration patterns (15–17).

For purposes of comparison, we map anatomical arbor size to the dimensionless load *A* using a simple proportional relationship,

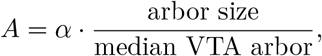

where *α* is chosen such that median SNc arbors fall near *A* ≈ 0.9–1.0. This mapping preserves **relative** differences between populations while acknowledging that a fully quantitative transformation from anatomy and physiology into (*A, C*) remains an open calibration problem. It is a provisional calibration for the present SNc/VTA contrast, not a prior-predictive estimate obtained independently of that contrast. **Because the bistable region in** (*A, C*) **space is relatively broad (Supplementary Figure S8), modest uncertainty in this mapping does not abolish the qualitative separation between VTA- and SNc-like parameter ranges; instead, it mainly shifts their positions along the diagonal bistable band**.

Taken together, the updated geometric picture provides a single unifying explanation:

- **VTA:** monostable → perturbations always recover.
- **Most SNc:** bistable → perturbations may push across the separatrix.
- **SNc outliers:** near right fold → collapse becomes nearly unavoidable.

Population heterogeneity in arbor size and Ca^2^ physiology enters the model exclusively through variation in (*A, C*). Even modest variability spreads neurons across the monostable and bistable regimes, naturally producing the observed heterogeneity in SNc vulnerability.

### 4.1 Population-level anatomical mapping of axonal load

To evaluate robustness to anatomical variability, we mapped synthetic populations of SNc and VTA neurons into *A*-space using literature-reported arborization statistics (1,3,15–17). Across 5,000 simulated neurons per group, VTA neurons formed a tight distribution centered around *A* ≈ 0.3–0.5, entirely outside the bistability window. SNc neurons formed a broader distribution centered near *A* ≈ 1.0, with the majority falling inside the bistable band (Supplementary Figure S12). Thus, within this provisional whole-cell mapping, realistic anatomical heterogeneity preserves the population-level segregation of VTA (stable) and SNc (susceptible) regimes.

This dynamic positioning—the placement of VTA and SNc neurons on opposite sides of a load-induced bistable window—provides a parsimonious explanation for selective vulnerability, contingent on the empirical mapping from anatomical and physiological load to the dimensionless parameters *A* and *C* and on the fact that the present formulation is spatially homogeneous rather than compartmentalized.

### 4.2 Approximate mapping from anatomy and physiology to (*A, C*)

Our dimensionless load parameters *A* and *C* are constructed to preserve empirically observed *ratios* between VTA and SNc neurons rather than to provide an absolute metric of metabolic burden. To make this mapping explicit, we start from literature estimates of (i) axonal arbor size and synapse number [1– 4,15–17] and (ii) Ca^2^-handling burden during pacemaking [5–9]. Single-neuron reconstructions suggest that SNc dopaminergic neurons form roughly 4–10× more synapses in the striatum than typical VTA neurons [1,3,15,16]. We therefore define a dimensionless arborization index

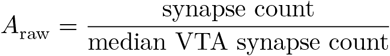

and then rescale by a constant factor *α* such that the empirical VTA range *A*_raw_ ≈ 1–1.5 maps to *A* ≈ 0.3–0.5 and the empirical SNc range *A*_raw_ ≈ 4–8 maps to *A* ≈ 0.9–1.0. This choice ensures that the observed 4–8× SNc:VTA synapse ratio corresponds to a approximately 2–3× increase in effective axonal load within the model, reflecting the fact that some components of baseline energy consumption (e.g., housekeeping costs) do not scale linearly with synapse number [24,25].

A similar construction is used for Ca^2^-handling load. Electrophysiological and imaging data indicate that SNc neurons experience greater L-type Ca^2^ influx during pacemaking than many VTA neurons [5–9]. We define a raw Ca^2^ load index proportional to the time-averaged Ca^2^ current during spontaneous firing and normalize so that typical VTA values correspond to *C* ≈ 1 and SNc values lie modestly above this range. Because *A* and *C* enter the equations only through their product *AC*, modest uncertainty in either component mainly shifts populations along the diagonal bistable band in (*A, C*) space rather than moving them in or out of the band (Supplementary Figure S8). We therefore interpret (*A, C*) as *effective* loads whose exact numerical values are uncertain but whose relative ordering between SNc and VTA is empirically grounded.

Importantly, the bistable band in (*A, C*) space is wide enough that realistic errors in this mapping do not collapse the separation between SNc-like and VTA-like regimes. VTA neurons remain in the monostable region for a broad swath of plausible (*A, C*) values, while SNc neurons—with their much larger arbors and higher Ca^2^ burden—robustly fall within or near the bistable band [1,3,5–9,15–17].

## 5. Perturbation Experiments Demonstrate Collapse in SNc but Recovery in VTA Neurons

The updated bifurcation structure provides a mechanistic foundation for interpreting the perturbation experiments. Because **VTA neurons lie entirely left of the left fold (***A* < 0.86), they possess **only one attracting state**, and any perturbation eventually returns to that high-energy equilibrium.

In contrast, **SNc neurons (***A* ≈ 0.9**–**1.0) lie **inside the bistable window**, where the saddle’s stable manifold divides the high- and low-energy basins. Perturbations that move the trajectory across this manifold, even transiently, inevitably carry the system to the collapsed low-energy attractor.

The perturbation experiment therefore operationalizes the bifurcation structure:

- For **VTA** (*A* = 0.40): the system returns to the high-energy attractor because no competing attractor exists.
- For **SNc** (*A* = 1.00): the same perturbation shifts the trajectory into the collapsed basin, producing an irreversible decline.

To focus on the geometric consequences of energy depletion, we implemented a minimal perturbation protocol that directly resets the energy variable while leaving mitochondrial capacity unchanged. In biological settings, insults often affect both *E* and *M*; because the saddle’s separatrix is codimension one, any joint perturbation that crosses the basin boundary has the same qualitative consequence. We therefore use the energy-only perturbation in the main simulations to isolate the geometric effect of transient energetic depletion, and examine joint (*E, M*) perturbations in Supplementary Figure S10.

To test how each neuronal population responds to transient metabolic stress, we simulated energy trajectories beginning near the high-energy steady state for both load conditions (*A* = 0.40 for VTA-like neurons and *A* = 1.00 for SNc-like neurons). Under these baseline conditions, both cell types remain stable and maintain high energetic reserve over long timescales (Figure 5A). This confirms that elevated structural load alone does not force SNc neurons into the pathological state; rather, it places them near a boundary where recovery from perturbation becomes precarious (15–17,24).

**Figure 5.**
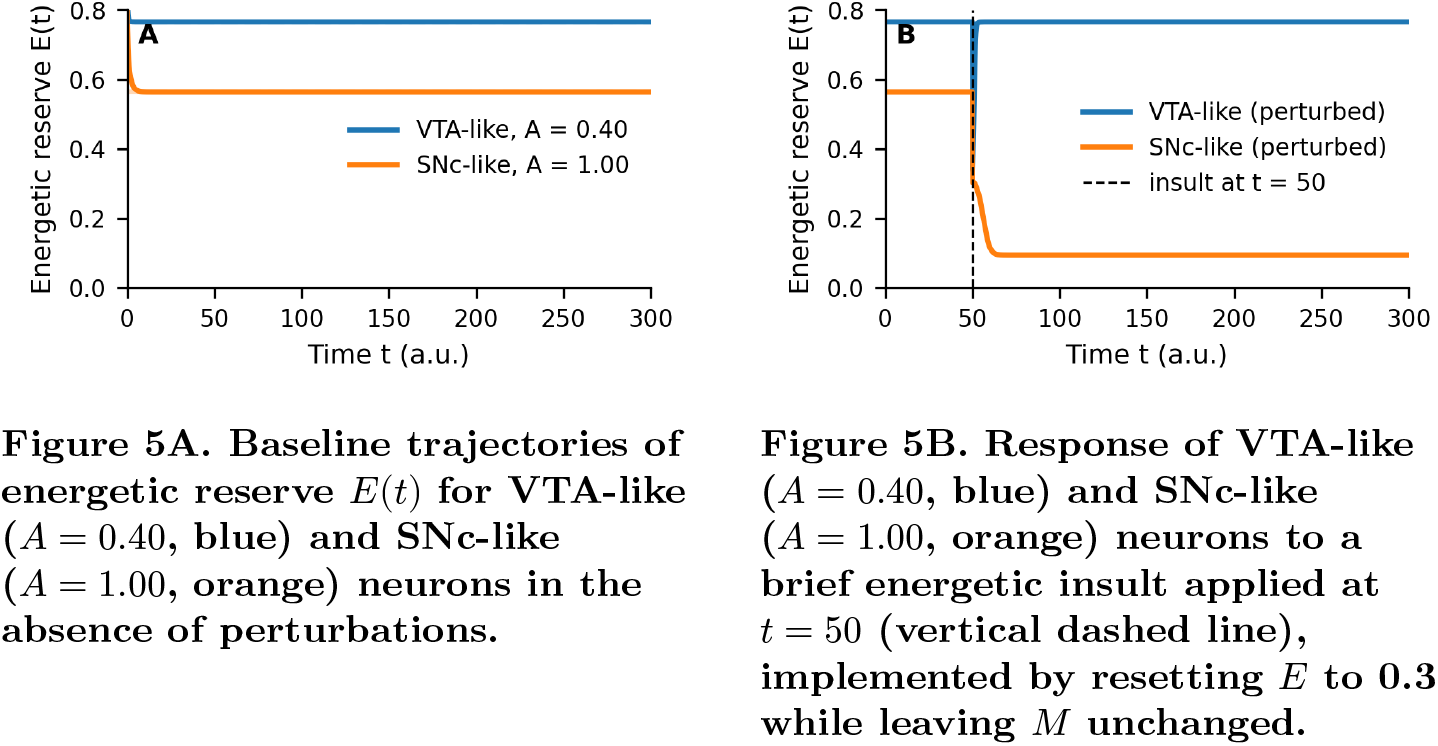
Time-course simulations reveal robustness in VTA neurons and collapse in SNc neurons. *Left (Figure 5A):* Starting from slightly elevated initial conditions, both traces relax rapidly onto their respective high-energy steady states (dashed lines) and remain stable over long times, while the corresponding mitochondrial capacities *M*(*t*) (not shown) behave similarly. Elevated structural load lowers the SNc steady-state energy level but does not drive spontaneous decline, indicating that extreme arborization alone is compatible with a persistent high-energy operating point and that collapse requires an additional perturbation. *Right (Figure 5B):* In the VTA-like monostable regime, *E*(*t*) rapidly relaxes back to its original high-energy steady state, indicating robust recovery from transient metabolic stress. In the SNc-like bistable regime, the same perturbation pushes the system across the saddle’s separatrix so that *E*(*t*) subsequently drifts toward and stabilizes at the low-energy attractor; the corresponding mitochondrial capacity *M*(*t*) (not shown) declines in parallel. The divergent outcomes of identical energy-only insults in the two regimes illustrate how proximity to a load-induced tipping point allows SNc neurons—but not VTA neurons—to undergo irreversible energetic collapse after modest perturbations (15–17,21–24). The divergent outcomes reflect the underlying folds at *A* = 0.86 and *A* = 1.06: VTA-like neurons sit left of the bistable window; SNc-like neurons sit inside it, where perturbations can push trajectories across the separatrix.

**Figure 6.**
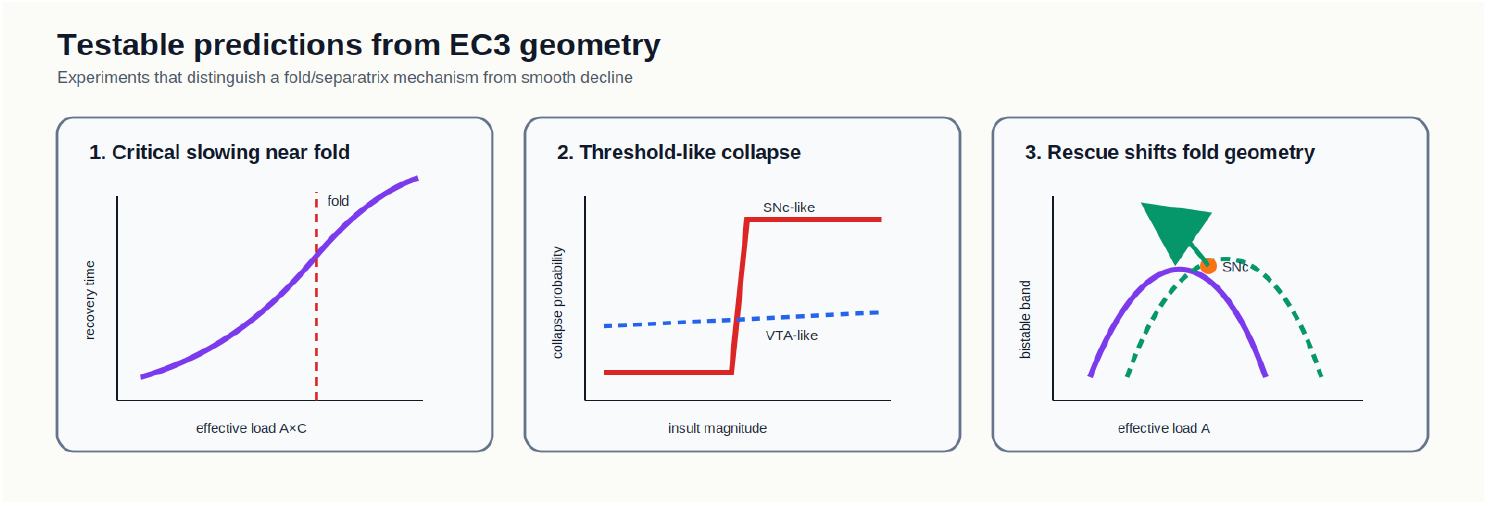
Testable predictions from EC3 geometry. The model predicts three experimentally distinguishable signatures of a fold/separatrix mechanism. First, recovery from small perturbations should slow as effective load approaches a fold. Second, SNc-like cells near the separatrix should show threshold-like collapse as insult magnitude increases, whereas VTA-like cells should recover across the same range. Third, interventions that reduce Ca^2^-handling burden, improve mitochondrial turnover, or reduce load-dependent damage should shift the fold geometry and move SNc-like neurons away from the separatrix.

We introduced a brief energetic perturbation by transiently reducing the energy variable *E* to 0.3 at time *t* = 50, mimicking a short-lived metabolic challenge such as a burst of pacemaking Ca^2^ entry, local inflammation, oxidative stress, or a mitochondrial inhibition event (5–9,10–14,24). The subsequent trajectories reveal a marked divergence between the two neuronal types (Figure 5B).

In the **VTA-like regime** (*A* = 0.40), the system quickly returns to the high-energy steady state after the perturbation. The energy reserve recovers smoothly, and mitochondrial capacity stabilizes along the same trajectory as in the unperturbed baseline. This behavior reflects the fact that VTA neurons, with their modest arborization load, lie far from the saddle-node bifurcation and thus possess a **single, globally attracting energetic state**. Perturbations may transiently reduce energy but do not threaten long-term stability (15–17).

In contrast, the **SNc-like regime** (*A* = 1.00) shows a dramatically different response. The same perturbation pushes the system across the stable manifold of the saddle, causing it to exit the basin of attraction of the high-energy state and converge instead to the low-energy attractor. Within the parameter range considered here, this collapse is effectively irreversible: simply returning to baseline loads does not restore the high-energy state because the trajectory remains in the collapsed basin. The resulting behavior reflects the hallmark of a **tipping-point transition**: the neuron appears stable until a modest perturbation triggers a sudden and catastrophic drop in energetic reserve from which recovery is no longer possible (21–23).

To verify that collapse is not specific to uni-dimensional perturbations of *E*, we also examined joint perturbations in (*E, M*) space near the SNc-like equilibrium (Supplementary Figure S10). Perturbations slightly above the separatrix returned to the high-energy state, whereas those slightly below it fell into the low-energy basin. This confirms that switching between stable states is determined by the geometry of the separatrix rather than by the direction of perturbation: any multidimensional insult—energetic, excitotoxic, oxidative, or metabolic—that crosses this boundary drives the system to the low-energy state.

This behavior mirrors the clinical and pathological course of Parkinson’s disease, where dopaminergic neurons can maintain function for decades before undergoing a rapid and irreversible decline (15–17,24). The model suggests that this vulnerability arises not from uniquely weak mitochondria or exclusive molecular stressors but from the **geometry of the underlying energy–mitochondria feedback loop**. SNc neurons operate close to a separatrix due to their extreme anatomical and physiological load. VTA neurons, lacking this structural burden, remain comfortably within a monostable regime where perturbations do not precipitate collapse.

These perturbation experiments thus provide computational evidence that **proximity to a load-induced saddle-node bifurcation is sufficient to explain the observed differential response to energetic insults under the assumptions of this minimal model** (15–17,21–23). Exploring a broader range of insult modalities—including joint perturbations to *E* and *M* and more detailed biophysical perturbation models—will be important to assess how general these conclusions remain across more biologically detailed perturbations.

In this first analysis we treated the system as deterministic, ignoring intrinsic and extrinsic noise in metabolic processes. One might worry that if SNc neurons operate near a separatrix, modest stochastic fluctuations in *E* or *M* could themselves trigger frequent noise-induced transitions into the collapsed basin [21–23]. We note that crossing the separatrix requires coherent excursions in the (*E, M*) plane of a magnitude comparable to the perturbations applied in our simulations, whereas single-channel and mitochondrial noise typically produces much smaller fluctuations in membrane potential and ATP levels on short timescales [10–14,24,25]. A full treatment of stochasticity would augment the ODEs with noise terms and quantify first-passage times to the collapsed state as a function of (*A, C*) and noise amplitude. Such an analysis is beyond the scope of this minimal deterministic study, but conceptually the results would be straightforward to interpret: if physiologically realistic noise amplitudes are insufficient to cross the separatrix, then collapse requires explicit insults of the kind we model here; if noise alone is sufficient, then the saddle-node geometry would still govern which populations (SNc vs VTA) are susceptible, but transitions would occur stochastically rather than only in response to discrete insults. Either way, the presence or absence of a bistable window in (*A, C*) space remains the primary determinant of which neurons are vulnerable to noise-amplified collapse.

## 6. Discussion

The updated continuation analysis clarifies the central thesis: **selective SNc vulnerability is rooted in their location inside a load-induced bistable window**, bounded by saddle-node folds at

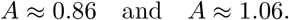

**The** (*A, C*) **stability map (Supplementary Figure S8) shows that these folds extend across Ca**^**2**^ **-handling loads as an oblique bistable band: neurons with high** *A* **and high** *C* **are pulled into the band, whereas neurons with low arborization and/or low Ca**^**2**^ **load remain in the monostable region**. Two-parameter sweeps across *A* and *k*_2_ further demonstrate that the bistable band persists over a broad range of nonlinear amplification strengths (Supplementary Figure S11). Together, these analyses show that the bistable energetic structure is preserved across joint variations in axonal load, Ca^2^-handling load, and nonlinear metabolic gain. This robustness indicates that the SNc’s vulnerability does not depend on fine parameter tuning, but reflects a geometric feature of the system’s load-balanced dynamics.

Under the representative calibration used here, **VTA neurons** (*A* ≈ 0.3–0.5) sit comfortably **left of the left fold**, in a monostable regime where energetic recovery is expected within the model. **SNc neurons** (*A* ≈ 0.9–1.0) sit **inside the bistable window**, close to the saddle separatrix. And **SNc outliers** (*A* ≈ 1.0–1.1) approach the **right fold**, where the high-energy state disappears entirely.

This geometry transforms the longstanding puzzle of selective vulnerability into a dynamical one:

- SNc neurons are not “weak”; they are operating on the interior of a bistable regime where **even modest perturbations** can initiate effectively irreversible collapse.
- VTA neurons are not “resilient” in absolute terms; they simply operate in a **monostable region** where collapse is structurally impossible within the model.
- Rescue by pruning in this framework requires reducing effective load below the left fold (*A* ≲ 0.86); in the absence of concurrent improvements in mitochondrial turnover or reduced calcium burden, such a reduction appears implausible for typical SNc neurons given current anatomical estimates.

Population-level simulations further demonstrate that realistic variability in arbor size does not blur regime separation: VTA neurons remain consistently outside the bistable window, whereas the majority of SNc neurons lie within it (Supplementary Figure S12). This reinforces the anatomical plausibility of the tipping-point mechanism at the population level.

The saddle-node geometry also makes a testable prediction: as parameter values approach a fold, the dominant recovery rate should slow, producing critical-slowing signatures in sufficiently resolved metabolic time series. A stochastic extension of the model confirms the same qualitative ordering: VTA-like loads remain protected because no collapsed attractor exists, whereas SNc-like loads show increasing noise-induced crossing probability as the high-energy basin shrinks near the right fold (Supplementary Figure S15). Calibrating absolute noise amplitudes, variance, autocorrelation, and flickering statistics from experimental data remains an important next step.

Because the vector field points inward along all boundaries of the normalized state space, the model never generates unphysical negative energy or negative mitochondrial capacity, and all trajectories remain within the physiological domain [0, 1]^2^ (Supplementary Figure S9).

This dynamical picture can account for the hallmark clinical sequence: decades of apparent stability followed by sudden decline. It also clarifies therapeutic leverage points: interventions that shift the system *leftward* (lower effective *A* or *C*), *upward* (improving mitochondrial turnover), or *rightward* (deepening the high-energy basin) all move the neuron further from the saddle and reduce collapse probability.

Our claim is explicitly one of **sufficiency**: a load-induced saddle-node geometry, instantiated in a minimal two-variable model, can reproduce key qualitative features of selective SNc vulnerability. This result does not establish uniqueness. Noise-induced escape from a monostable landscape, slow drift in parameters, threshold switching, or other compact mechanisms could also generate abrupt failure under some conditions. Distinguishing these alternatives will require matched model comparisons using common perturbation protocols, calibrated noise amplitudes, and dynamical summary statistics such as recovery rates, first-passage times, and basin-boundary estimates. The value of the present model is that the saddle-node motif arises directly from empirically observed structural and Ca^2^-handling loads and yields concrete, testable predictions about fold locations, separatrix crossing, and rescue by moving the system out of the bistable band.

The model is also complementary to existing computational accounts rather than a replacement for them. Prior work has quantified the energetic cost of action-potential propagation through large dopaminergic arbors and developed conductance-level models of dopamine-neuron pacemaking, bursting, and calcium dynamics (31,34–37). Those models operate at a more biophysical level and are natural sources for future calibration of the effective loads *A* and *C*. The present model deliberately compresses those details into a two-dimensional energetic phase plane to ask a different question: whether known structural and calcium-associated burdens are sufficient to place SNc-like neurons in a different dynamical regime than VTA-like neurons.

This framing yields several experimentally testable predictions: (i) metabolic recovery rates should slow as cells approach the fold; (ii) equal energetic insults should have threshold-like outcomes near SNc-like but not VTA-like loads; (iii) interventions that reduce Ca^2^-handling burden or improve mitochondrial turnover should shift the fold structure and enlarge the high-energy basin; and (iv) molecular insults such as dopamine oxidation, -synuclein toxicity, or mitophagy impairment should primarily act by moving the cell closer to the separatrix through changes in *M, k*_*M*_, *β*, or effective load.

This energetic-tipping framework is explicitly not a replacement for molecularly detailed accounts of Parkinson’s pathology. Rather, it provides a dynamical context into which mechanisms such as -synuclein aggregation, dopamine oxidation, impaired mitophagy, and LRRK2- or Parkin-mediated mitochondrial dysfunction can be naturally embedded [10–14,18–20]. In our model, such processes would primarily act by shifting parameters that modulate mitochondrial capacity and damage (e.g., *k*_*M*_, *β*) or effective load (via changes in firing rate and Ca^2^ influx), thereby moving neurons toward or away from the bistable band in (*A, C*) space. For example, -synuclein–induced disruption of mitochondrial dynamics would effectively lower *M* or *k*_*M*_, narrowing the high-energy basin at a given (*A, C*); increased oxidative stress from dopamine metabolism would increase the effective damage term *βACM*(1 − *E*); and calcium-channel “rejuvenation” strategies [7–9] can be interpreted as reducing the co-load *C* and thus moving SNc neurons leftward in (*A, C*) space, away from the bistable window. On this view, molecular insults do not create SNc vulnerability de novo; they determine how close structurally heavily loaded neurons are to the saddle-node folds, and therefore how readily insults or noise can push them across the separatrix.

Our joint (*E, M*) perturbation experiments (Supplementary Figure S10) show that collapse is inherently geometric: any multidimensional insult that crosses the separatrix—energetic, excitotoxic, oxidative, or metabolic—drives the system to the low-energy state. The stochastic analysis (Supplementary Figure S15) extends this result from imposed insults to fluctuation-driven escape, showing that collapse probability depends strongly on load-dependent basin geometry and noise amplitude. This provides a unifying interpretation for diverse experimentally observed injury modalities. Numerical robustness analyses across solver families and tolerances (Supplementary Figure S13) further confirm that the classification of trajectories into high-energy versus low-energy asymptotic states is independent of solver choice and tolerance within a broad range, excluding the possibility that the bistable structure is a numerical artifact.

The signs and qualitative magnitudes of all model terms correspond to well-established empirical relationships: increased arbor size raises ATP demand; Ca^2^ entry elevates mitochondrial load; energetic deficit impairs mitochondrial repair; and preserved mitochondrial capacity supports ATP production (5–14,24,25). Although a precise quantitative calibration is not yet available, the **directionality** of these interactions is directly grounded in experimental literature.

Because the model is presently nondimensional in time, we have not attempted to fit absolute progression timescales. Our claims are therefore geometric rather than temporal: we show that realistic structural and physiological loads can place SNc neurons inside a bistable energetic regime while VTA neurons remain outside it. Calibrating the model so that trajectories reproduce decade-long prodromal periods and age-dependent incidence curves is an important next step that will require quantitative data on mitochondrial turnover and energy budgets specific to SNc and VTA neurons.

Finally, our regime assignments for SNc and VTA neurons rely on **provisional calibration** of the dimensionless load parameters. We map *A* onto relative differences in arbor size and synapse number from anatomical reconstructions and use *C* to capture Ca^2^-handling burden, but we do not yet possess a fully quantitative transformation from measured morphology and physiology into (*A, C*) with uncertainty bounds. We therefore interpret our population-level statements probabilistically: most SNc neurons are predicted to occupy the bistable regime and most VTA neurons the monostable regime, given current anatomical and physiological evidence, and we explicitly highlight the sensitivity of these conclusions to the load mapping, **even though the** (*A, C*) **bistable band itself is wide enough that moderate calibration errors do not collapse the separation (Supplementary Figure S8)**.

The right fold at *A* ≈ 1.06 exceeds known anatomical loads for most SNc neurons and is therefore primarily of mathematical interest. Physiologically, neurons operate well to the left of this second fold, and the **left fold** is the barrier that determines whether recovery to a healthy state is dynamically permissible. Within this class of models, collapse at typical SNc loads cannot be undone simply by returning to baseline anatomy; instead, one must move the neuron out of the bistable regime altogether or modify other parameters that reshape the folds.

The model therefore offers a unifying principle: **extreme combined structural and calcium-handling load can place SNc neurons in a fundamentally different dynamical regime than VTA neurons**. Their vulnerability follows from the geometry of this class of models, rather than requiring a cell-type-specific molecular anomaly—provided that physiological parameter values indeed place them inside the bistable window implied by the feedback motif.

### Limitations and future directions

Several limitations of the present model deserve emphasis. First, we consider only two state variables (*E* and *M*) and treat axonal arborization and Ca^2^-handling load as static parameters (*A, C*), rather than modeling activity-dependent remodeling or structural plasticity. Incorporating slow dynamics for *A* and *C* could capture feedback between disease progression, synaptic loss, and firing patterns. Second, we omit explicit representations of -synuclein aggregation, dopamine oxidation, reactive oxygen species, and autophagic/lysosomal pathways, all of which are known to contribute to dopaminergic neuron degeneration [10–14,18–20]. As argued above, these mechanisms can be naturally incorporated as modulators of *k*_*M*_, *β*, and effective load, but doing so will require quantitative data to constrain how they reshape the folds in (*A, C*) space. Third, we work with nondimensional time and parameters and therefore do not yet reproduce absolute disease timescales. Likewise, we have not yet carried out a prior-predictive or Bayesian calibration of (*A, C, k*_2_) under independent priors, so the probability that the SNc/VTA separation appears without outcome-conditioned tuning remains unknown. A blind leave-one-population-out stress test reduced but did not eliminate circularity, so we retain it as a blocker note rather than a standalone claim. Fourth, a minimal two-compartment soma/distal-arbor extension with linear exchange coupling preserves a bistable whole-cell separatrix in finite load windows, but it also shifts the fold locations and makes distal-local perturbations more effective than soma-local ones. In this extension the bistable band is split by a monostable gap, suggesting that spatial compartmentalization modulates, rather than destroys, the tipping geometry. Fifth, the current mapping from morphology and Ca^2^ physiology to (*A, C*) remains provisional and was chosen for the SNc/VTA comparison rather than fit independently of it. Finally, the stochastic analysis is intentionally dimensionless: it quantifies how noise-induced separatrix crossing scales with load and basin geometry, but calibrated experimental time series will be required to map the noise amplitude *σ* to physiological ATP or mitochondrial fluctuations. A clinically relevant comparison is isradipine/STEADY-PD III: in this geometry, lowering *C* should shift trajectories leftward, but a negative trial is still compatible with an insufficient effective shift, intervention after separatrix crossing, or compensatory homeostatic responses that offset the calcium-load reduction. More broadly, the model is conceptually compatible with bounded-clearance aggregation frameworks, in which finite removal capacity generates hysteresis; coupling … [truncated]

## 7. Methods

This section describes the mathematical model, parameterization, numerical procedures, and perturbation protocols used in all analyses. The goal is transparency and reproducibility rather than biological completeness: the model is intentionally minimal, and all simulations use the same governing equations without case-specific modifications.

### 7.1 Model Equations

The energetic state of a dopaminergic neuron is represented by two normalized variables:

- **Energetic reserve:** (E(t) [0,1]), reflecting ATP/NADH availability.
- **Mitochondrial functional capacity:** (M(t) [0,1]), representing the effective ability of mitochondria to sustain oxidative phosphorylation.

The governing equations are:

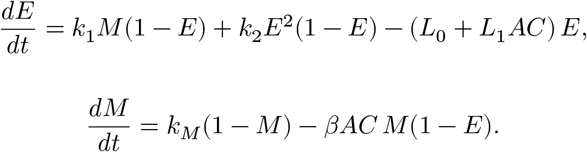

Two dimensionless load parameters modulate demand and damage:

- *A*: axonal arborization load.
- *C*: Ca^2^-handling load.

Unless stated otherwise, *C* = 1. All analyses interpret results within the physiological state domain [0, 1]^2^.

### 7.2 Parameter Values

All core simulations use the canonical EC3 parameter set:

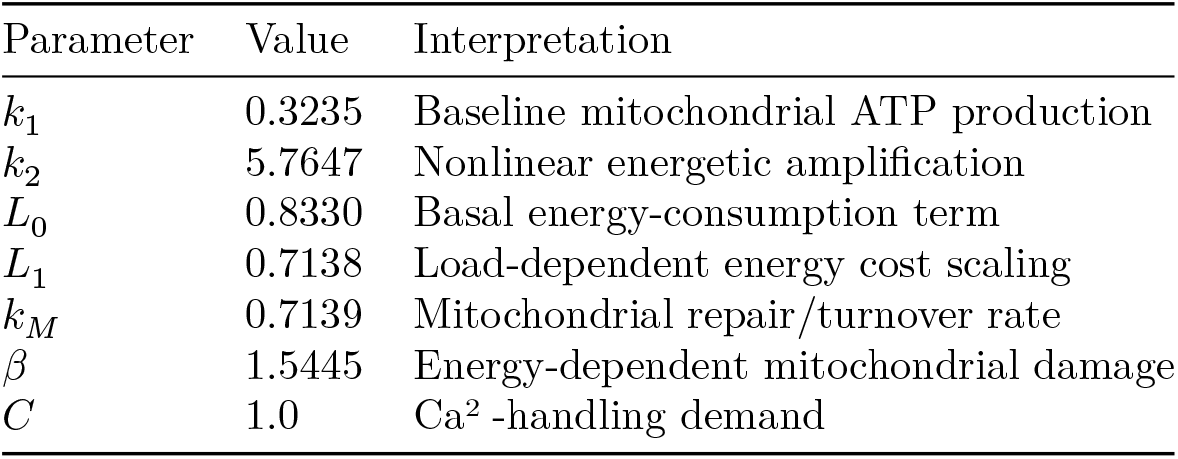

Values are rounded for readability; exact values are provided in the repository. Because the state variables are normalized and the time variable is nondimensional, the listed parameters should be read as dimensionless rates rather than physical constants with SI units.

These parameters are phenomenological and nondimensional rather than direct fits to a single experiment or a prior-predictive calibration against independent data. Their provenance and intended interpretation are:

| Quantity | Biological interpretation | Current provenance / uncertainty |
| --- | --- | --- |
| $E$ | Energetic reserve, integrating ATP/ADP balance, NADH/FAD state, spare respiratory capacity, and ability to meet acute demand | Coarse normalized state variable; potential experimental proxies include ATP/ADP ratio, NADH/FAD autofluorescence, mitochondrial membrane potential, and oxygen consumption (24,25,38–41) |
| $M$ | Functional mitochondrial capacity, including oxidative phosphorylation competence, repair, turnover, and stress-induced dysfunction | Coarse normalized state variable; potential proxies include respiratory capacity, membrane potential, mitophagy/turnover markers, and mitochondrial oxidant stress (10–14,33,42) |
| $A$ | Effective axonal structural load | Direction constrained by single-axon/arborization studies and energetic-cost modeling, but the mapping from measured arbor size to dimensionless $A$ remains provisional (1,3,15,29–32) |
| $C$ | Effective $\text{Ca}^{2+}$-handling/metabolic co-load from pacemaking and calcium buffering | Direction constrained by pacemaking/$\text{Ca}^{2+}$ studies; treated here as a sensitivity parameter rather than a calibrated conductance-level variable (5–9,33–37) |
| Rate parameters | $k_1, k_2, L_0, L_1, k_M$, and $\beta$ are normalized rates controlling ATP support, nonlinear energetic amplification, consumption, mitochondrial repair, and load-dependent damage | Chosen to instantiate a representative EC3 regime; robustness is assessed by parameter sweeps, but empirical fitting remains future work |

For population comparisons:

- **VTA-like:** *A* = 0.40
- **SNc-like:** *A* = 1.00

These assignments map literature-reported relative arbor sizes onto the dimensionless load scale and place each population on the appropriate side of the bifurcation structure.

### 7.3 Initial Conditions and Perturbation Protocol

All simulations begin at the stable high-energy equilibrium (*E*^*^, *M*^*^) corresponding to each load (*A, C*). This reflects the fact that dopaminergic neurons typically operate near a long-lived energetic steady state.

#### Energetic perturbations

To assess vulnerability to transient metabolic insults, we applied fast-timescale perturbations implemented as instantaneous reductions of energetic reserve:

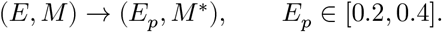

This captures biologically common events—ATP depletion, mitochondrial inhibition, oxidative bursts—that reduce energetic reserve much more rapidly than mitochondrial repair/damage processes can respond. After the instantaneous reset, the system is integrated forward under the same ODEs, so both *E* and *M* continue evolving dynamically toward recovery or collapse; *E* is not held fixed after the insult. Because the saddle’s stable manifold is codimension-one, the direction of perturbation is irrelevant: any trajectory displaced across the separatrix inevitably falls into the low-energy basin, and any trajectory displaced within the healthy basin returns to the high-energy state.

#### Joint (*E, M*) perturbations

To ensure that outcomes were not specific to the *E*-only insult design, we also evaluated joint perturbations:

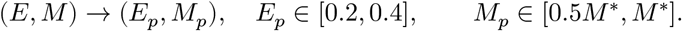

These represent insults that simultaneously impair energetic reserve and mitochondrial capacity (e.g., Ca^2^-mediated mitochondrial depolarization, oxidative mitochondrial damage). As shown in Supplementary Figure S10, perturbations on one side of the separatrix always recovered; those on the other side always collapsed. The geometric boundary, not the perturbation direction, determines the outcome.

#### Summary

- *E*-only perturbations probe the fast energetic timescale relevant for separatrix-crossing events.
- Joint perturbations confirm direction-independence of basin transitions.
- All conclusions about SNc vs VTA resilience arise from geometry, not perturbation design.

#### 7.4 Numerical Integration

All time-course simulations were performed using Python with SciPy for ODE integration, NumPy for numerical arrays, and Matplotlib for plotting (26–28). Exact package versions are recorded in the repository environment and can be reproduced from `requirements.txt` and the checked-in scripts.

We used SciPy’s explicit Runge–Kutta method of order 5(4) (**RK45**) (26) with default tolerances and a maximum step size of 0.5. Solutions were evaluated on 2000 uniform time points over *t* ∈ [0, 300].

#### Solver robustness

To confirm that collapse vs recovery classification was not a numerical artifact, we repeated all VTA-like and SNc-like perturbation simulations using:

- RK45 (explicit),
- RK23 (explicit, lower order),
- BDF (implicit, stiff),

each under default tolerances, 10× tighter tolerances, and 10× looser tolerances (9 total configurations). All solvers produced identical attractor outcomes with no spurious equilibria or oscillations (Supplementary Figure S13).

### 7.5 Equilibrium Identification and Stability Classification

For bifurcation analyses we located equilibria using a two-stage procedure:

1. **Coarse scan:** evaluate ODE right-hand sides on a grid in (*E, M*) to identify approximate zero-crossings.
2. **Refinement:** apply SciPy’s `root` (hybrid method) (26) to each candidate point, followed by deduplication within a Euclidean tolerance of 10^−3^.

Stability was determined by computing the Jacobian matrix at each equilibrium and examining eigenvalues:

- All eigenvalues with negative real parts → **stable** fixed point.
- One eigenvalue with positive real part → **saddle**.

The bistable window is defined as the set of *A* values, or (*A, C*) pairs, for which exactly **three** equilibria (two stable, one saddle) exist in [0, 1]^2^.

### 7.6 Bifurcation Scan Procedure

To construct the main one-parameter bifurcation diagram (Figure 3), axonal load was varied over:

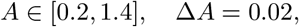

yielding 61 conditions. Each load was analyzed using the equilibrium-finding procedure in Section 7.5.

As an additional fold-location check, we solved the augmented system *dE*/*dt* = 0, *dM*/*dt* = 0, and det(*J*) = 0, where *J* is the local Jacobian. This refinement gives folds at approximately *A* = 0.849 and *A* = 1.075 for *C* = 1, consistent with the rounded values reported in the main text and figures.

For the extended continuation (Supplementary Figure S7), the range was expanded to *A* ∈ [0.2, 2.0] over 90 samples.

### 7.7 Phase-Plane Construction and Nullclines

Phase planes (Figures 2 and 4) were computed over a 25 × 25 grid in the domain [0, 1]^2^. At each grid point:

- the vector field (*dE*/*dt, dM*/*dt*) was evaluated and normalized for visualization;
- nullclines were traced using zero-contours of *dE*/*dt* and *dM*/*dt*;
- sample trajectories were generated via `solve_ivp` (26) from diverse initial conditions.

All computations were performed using the same governing equations and parameters as in Sections 7.1–7.2.

### 7.8 Stability Map in (*A, C*) Space

To characterize how axonal load and Ca^2^-handling load jointly govern dynamic regime boundaries, we computed a dense stability map on:

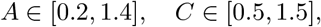

using a 121 × 121 grid. For each pair, the number of equilibria was determined as in Section 7.5. Regions with a single equilibrium were labeled **monostable**, and regions with three equilibria **bistable**. The resulting bistable band (Supplementary Figure S8) demonstrates the diagonal geometry of load-induced tipping across the (*A, C*) plane.

### 7.9 Two-Parameter Sweeps and Model Robustness

To assess robustness to uncertainty in nonlinear energetic amplification, we computed an additional stability map in (*A, k*_2_) space, sampling:

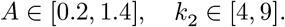

The bistable band persisted across the entire *k*_2_ range (Supplementary Figure S11), demonstrating that the saddle-node structure does not require fine-tuning.

### 7.10 Positive Invariance of the Physiological Domain

To ensure the model never produces unphysical negative energy or mitochondrial capacity, we evaluated the vector field along each boundary of [0, 1]^2^:

- At *E* = 0: *dE*/*dt* = *k*_1_*M* ≥ 0, with equality only at the corner *M* = 0.
- At *E* = 1: *dE*/*dt* < 0.
- At *M* = 0: *dM*/*dt* = *k*_*M*_ > 0.
- At *M* = 1: *dM*/*dt* ≤ 0 over the parameter ranges considered.

All vectors point inward (Supplementary Figure S9), confirming positive invariance: trajectories remain inside the physiological domain for all time.

### 7.11 Formal Dynamical Guarantees

The EC3 vector field admits several proof-level properties that are independent of the numerical scans.

Lemma 7.11.1 (Positive invariance of the physiological domain; cf. 21–23). The square [0, 1]^2^ is forward invariant for the EC3 flow.

Proof. The boundary signs are listed in Section 7.10: at *E* = 0, *dE*/*dt* = *k*_1_*M* ≥ 0; at *E* = 1, *dE*/*dt* = −(*L*_0_ + *L*_1_*AC*) ≤ 0; at *M* = 0, *dM*/*dt* = *k*_*M*_ > 0; and at *M* = 1, *dM*/*dt* = −*βAC*(1 − *E*) ≤ 0. Hence the vector field points inward (or is tangent at corners), so trajectories cannot exit [0, 1]^2^.

Lemma 7.11.2 (No oscillatory attractors in the interior; cf. 21,23,43). On (0, 1)^2^, the flow is cooperative: ∂*f*/∂*M* = *k*_1_(1 − *E*) > 0 and ∂*g*/∂*E* = *βACM* ≥ 0 (strictly positive whenever *A, C* > 0 and *M* > 0). Therefore the planar semiflow is strongly monotone in the interior, and standard monotone-dynamics results exclude periodic orbits or other non-equilibrium oscillatory attractors in the compact invariant domain.

Theorem 7.11.3 (Global attraction in the monostable regime; cf. 21,23,43). If, for a given (*A, C*), the reduced equilibrium equation *f*(*E*; *A, C*) = 0 has exactly one root in (0, 1), then that equilibrium is globally attracting in [0, 1]^2^.

Proof. By Lemma 7.11.1, every trajectory remains in the compact invariant set [0, 1]^2^, so every trajectory has a nonempty *ω*-limit set. By Lemma 7.11.2, that *ω*-limit set cannot contain a periodic orbit or any other oscillatory attractor. If the reduced equation has only one equilibrium in (0, 1), then that equilibrium is the only invariant limit set left in the domain, so every trajectory converges to it.

Theorem 7.11.4 (Persistence of bistability under saturating nonlinearities; cf. 21,23,44). Replace the polynomial cooperative term *k*_2_*E*^2^(1 − *E*) by a *C*^2^ saturating feedback *g*(*E*; *η*). If, for some base parameter value, the reduced equilibrium equation satisfies *f*(*E*; *A, C, η*) = 0 and ∂*f*/∂*E* = 0 at a point where ∂^2^*f*/∂*E*^2^ ≠ 0 and the unfolding direction satisfies ∂*f*/∂µ ≠ 0, then a nearby saddle-node persists under sufficiently small *C*^2^ perturbations of *g*.

Proof. The fold conditions define a codimension-one nondegenerate saddle-node in the reduced scalar equation. By the implicit function theorem, these conditions persist under sufficiently small *C*^2^ perturbations that preserve the local sign and curvature structure of the reduced nullcline, so a nearby fold branch exists for the perturbed saturating nonlinearity.

These statements formalize the three proof targets discussed in the text: positive invariance, absence of oscillations, and local persistence of the bistable geometry under broad saturating substitutions (cf. 43,44).

### 7.12 Stochastic Separatrix-Crossing Analysis

To quantify fluctuation-driven collapse without introducing new biological data, we simulated a dimensionless stochastic extension of the EC3 system:

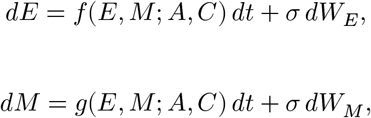

where *f* and *g* are the deterministic right-hand sides in Section 7.1 and *dW*_*E*_, *dW*_*M*_ are independent Wiener increments. Simulations used Euler–Maruyama integration with *dt* = 0.05 over *t* ∈ [0, 300], initialized at the high-energy equilibrium for each *A*. Boundary reflections kept trajectories inside the physiological domain [0, 1]^2^.

Because the present analysis is intended as a geometric sensitivity test rather than a calibrated noise model, *σ* was swept over dimensionless amplitudes. In bistable regimes, a stochastic trajectory was counted as collapsed when it remained on the low-energy side of the saddle’s *E* coordinate for at least two time units; outside the bistable regime, irreversible fluctuation-driven collapse was assigned zero probability because no collapsed attractor exists. For each (*A, σ*) pair, 220–250 stochastic paths were simulated and summarized by collapse probability and mean first-passage time. We also computed the minimum instantaneous energy drop from the high-energy equilibrium that deterministically enters the low-energy basin, plus the dominant linear recovery timescale at the high-energy equilibrium. Panel B lifts the zero-collapse SNc-edge curve slightly for visibility, and panel D uses color-matched dual y-axes and compact in-plot labels to distinguish the critical *E*-drop curve from the recovery-timescale curve. These analyses generate Supplementary Figure S15 and the accompanying CSV summary.

### 7.13 Reproducibility and Code Availability

Most simulations are deterministic; Supplementary Figures S15–S19 use fixed random seeds for reproducible stochastic/uncertainty trajectories. Full code— including ODE definitions, parameter sets, equilibrium scanners, bifurcation scripts, stochastic simulations, and figure-generation utilities—is available at:

**https://github.com/MaxAnfilofyev/parkinsons-ec3-model**

Running `python code/run_ec3_all.py` reproduces the main computed figures in the manuscript; the supplementary scripts listed in `docs/supplement.md` regenerate the supplementary figure set.

## Supplementary Results

### S1. Earlier model architectures did not exhibit bistability

We evaluated several alternative formulations of the energy–mitochondria interaction prior to arriving at the final minimal model. These earlier systems were structurally incapable of producing multiple equilibria under biologically realistic parameter ranges, despite including many of the same biological ingredients. Their failure highlights the importance of the nonlinear energy-amplification and load-modulated mitochondrial damage terms used in the final formulation (21–23).

#### S1.1 First model variant: linear mitochondrial support and linear load

The simplest architecture consisted of:

- Linear ATP production proportional to *M*(1 − *E*),
- Quadratic mitochondrial damage proportional to *ACM*(1 − *E*),
- No nonlinear amplification in energy (i.e., the *E*^2^(1 −*E*) term was absent).

In this system, the energy nullcline is monotonic for all parameter values, and the mitochondrial nullcline intersects it **exactly once** within the biologically relevant domain. The vector field analysis confirmed global convergence to a unique fixed point. No choice of axonal load *A*, even when increased far beyond anatomical ranges, produced a second intersection.

**Supplementary Figure S1.**
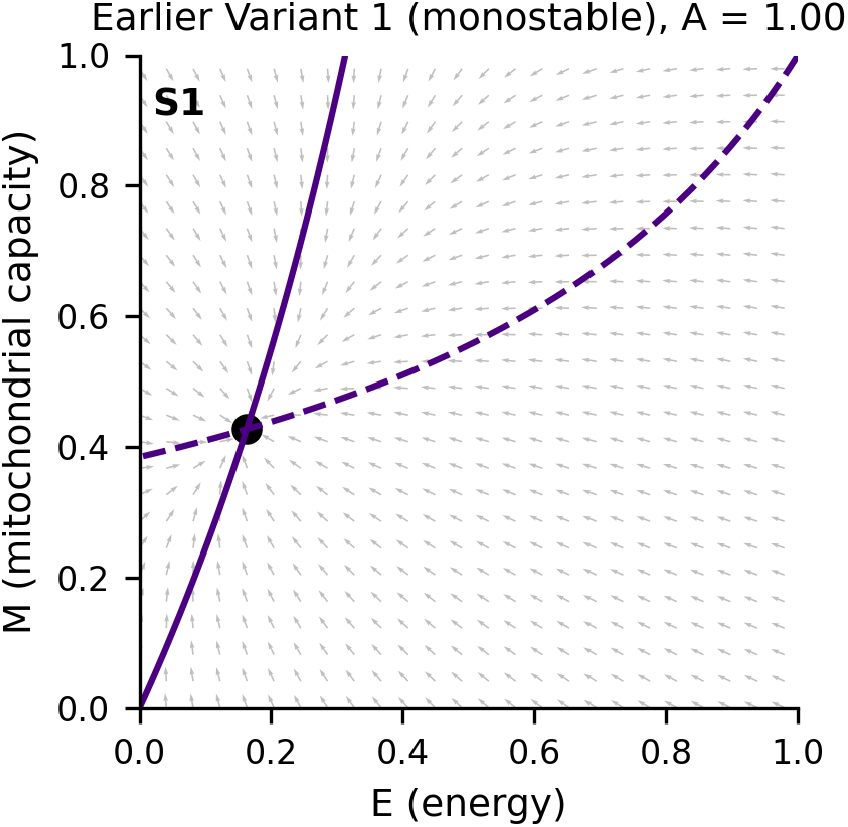
Nullclines and vector field for an earlier, monostable model variant. In a model with linear mitochondrial support and load effects but no nonlinear energy amplification, the energy and mitochondrial nullclines intersect only once in the physical domain. The vector field indicates global convergence to a single equilibrium for all tested values of *A*, demonstrating that this architecture cannot produce bistability.

#### S1.2 Second model variant: feedback mitochondrial impairment without nonlinear energy restoration

The second architecture attempted to incorporate a biologically motivated feedback: energetic deficit increases mitochondrial damage. Mathematically, this introduced a term of the form (1−*M*)*E* in the energy equation and retained the load-dependent mitochondrial damage. Despite this additional coupling, the system remained **monostable**.

- The energy nullcline remained single-peaked but never developed a fold.
- The mitochondrial nullcline cut across it only once.
- Bifurcation scans across a wide range of *A* ∈ [0.1, 2.0] showed one stable equilibrium everywhere.

**Supplementary Figure S2.**
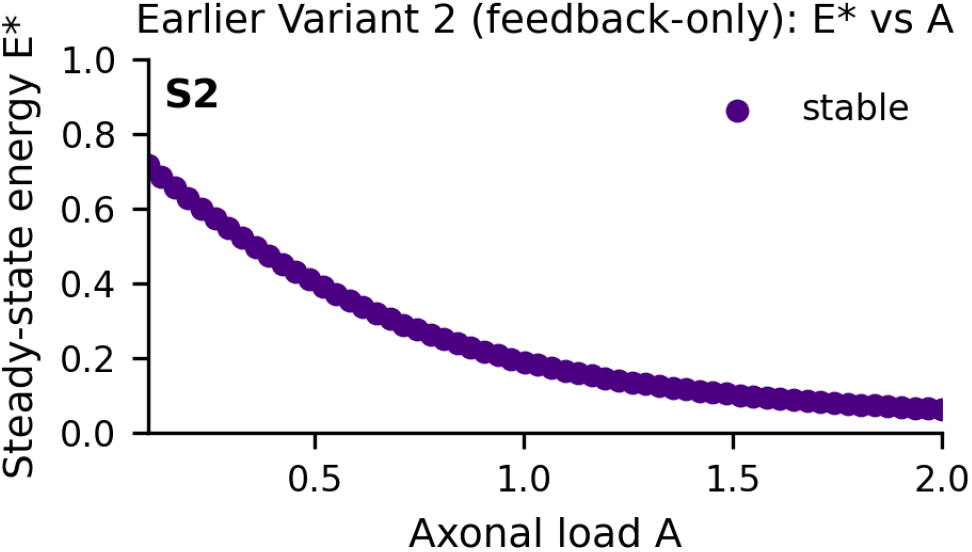
Bifurcation scan for a feedback-only model lacking nonlinear energy amplification. Even with feedback from energy to mitochondrial damage, the system exhibits only a single equilibrium across a broad range of axonal loads *A*. No saddle-node bifurcation appears, confirming that feedback alone is insufficient to generate bistability in this architecture (21–23).

These results demonstrated that feedback alone is insufficient: to generate bistability, the model must produce a genuine **S-shaped energy nullcline** within the physical domain.

### S2. Requirement for nonlinear energy amplification

The term *k*_2_*E*^2^(1 − *E*) in the final model introduces a saturating, cooperative-like energetic contribution that is negligible near *E* = 0 but increases sharply as *E* rises. This positive curvature is what allows the energy nullcline to fold under load (21–23).

By contrast:

- In models lacking this term, the nullcline was monotonic.
- In models using only linear or quadratic forms, the nullcline never turned sufficiently to create two additional equilibria.

When included, the nonlinear term interacts with load-dependent consumption and energy-dependent mitochondrial damage to produce:

1. A high-energy fixed point,
2. A low-energy fixed point,
3. A saddle separating their basins.

**Supplementary Figure S3.**
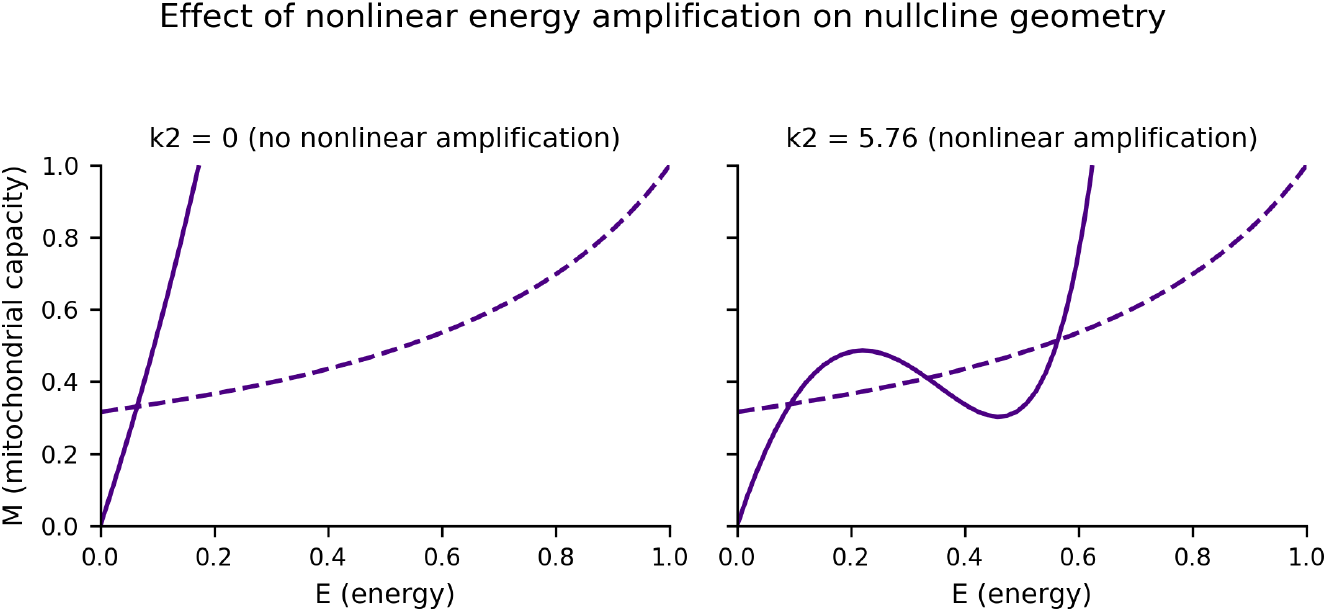
Effect of nonlinear energy amplification on nullcline geometry. Including the nonlinear term *k*_2_*E*^2^(1 − *E*) generates a folded energy nullcline that, together with the mitochondrial nullcline, produces three intersections corresponding to two stable equilibria and a saddle. Removing this term collapses the fold and eliminates bistability.

### S3. Parameter sweeps confirm robustness of the bistable window

We systematically varied key parameters to assess how sensitive the saddle-node structure is to physiological uncertainty (Supplementary Figures S4–S6). In keeping with the minimal and analytically transparent nature of the model, we focused on one-at-a-time parameter sweeps rather than fully joint sampling across all parameters.

### S3.1 Variation in mitochondrial turnover rate *k*_*M*_

**Supplementary Figure S4.**
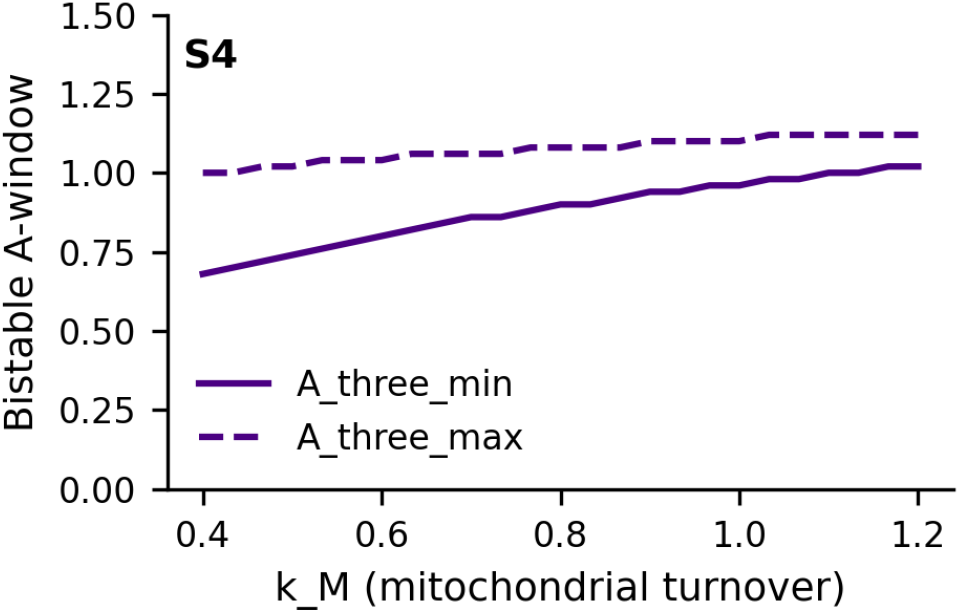
Effect of mitochondrial turnover rate *k*_*M*_ on the bistable window in axonal load *A*. Increasing *k*_*M*_ shifts the bistable window to higher values of *A*, reflecting improved mitochondrial resilience. Conversely, reducing *k*_*M*_ expands the bistable region toward lower loads, making more neurons susceptible to collapse (10–14,21–23).

### S3.2 Variation in load-dependent consumption *L*_1_

**Supplementary Figure S5.**
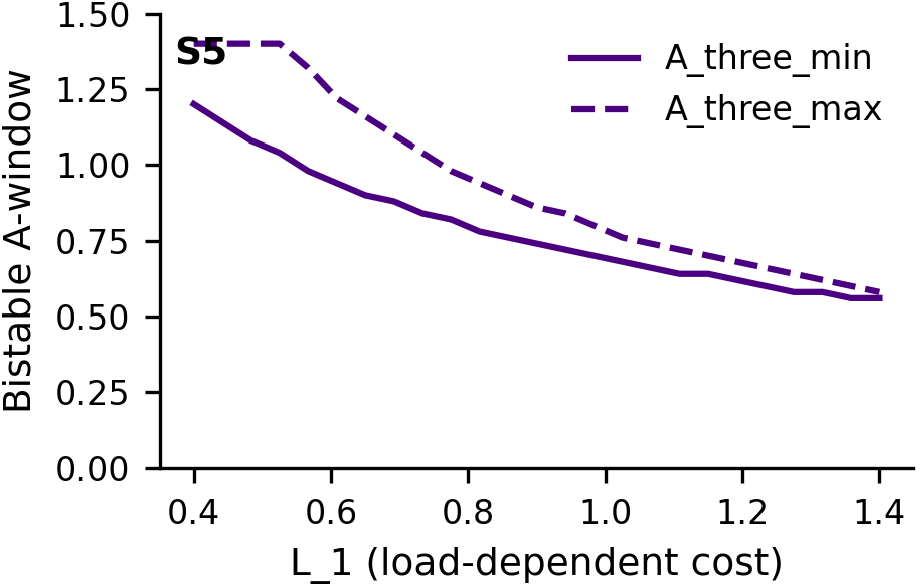
Effect of load-dependent energy cost *L*_1_ on the bistable window in axonal load *A*. Higher *L*_1_ (greater energy cost per unit arbor) shifts the bistable window to lower values of *A* and lowers the healthy branch; lower *L*_1_ shifts the window upward and raises the healthy equilibrium.

### S3.3 Variation in calcium-handling load *C*

**Supplementary Figure S6.**
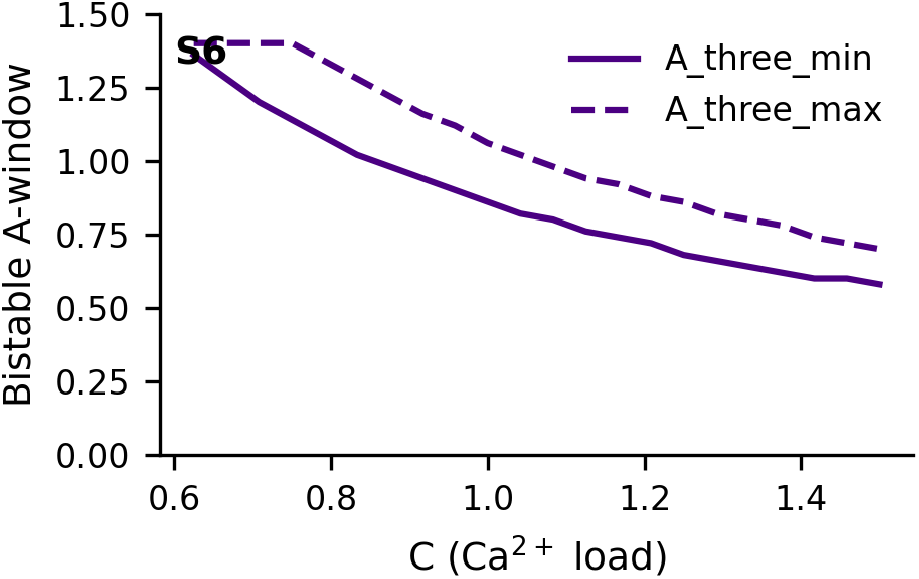
Effect of Ca^2^-handling load *C* on the bistable window in axonal load *A*. Because SNc neurons experience substantial Ca^2^ influx during pacemaking, we explored *C* ∈ [0.5, 1.5]. Changes in *C* functionally act like scaling *A*; high *C* shifts the bistable window leftward, increasing vulnerability (5–9,21–23). **To go beyond one-dimensional sweeps, we computed a high-resolution stability map across the full** (*A, C*) **plane (Supplementary Figure S8). This analysis confirms that the bistable region forms a continuous band: increasing** *C* **shifts the bistable window to lower** *A*, **decreasing** *C* **shifts it to higher** *A*, **and there is no isolated island of bistability**.

Across these one-at-a-time sweeps, the saddle-node bifurcation persisted, demonstrating that the fold emerges robustly from the feedback motif under a broad range of individual parameter variations. Assessing robustness under correlated multi-parameter variation—for example, jointly varying *k*_*M*_ and *C*—is an important next step for quantifying how sensitive the bistable window is to combined physiological uncertainty. The (*A, C*) stability map in Supplementary Figure S8 provides a compact geometric summary of these effects.

### S4. Absence of biologically relevant hysteresis

The updated full continuation analysis (Figure 3) reveals that the saddle-node bifurcation contains two folds:

- A **left fold** at *A* ≈ 0.86, where bistability first appears.
- A **right fold** at *A* ≈ 1.06, where the high-energy and saddle branches merge and annihilate.

Because **typical SNc loads** lie in *A* ≈ 0.9–1.0, they are **inside** the bistable window but safely **left of** the right fold.

This implies:

1. **Collapse is not recoverable by returning to typical SNc loads**. Once a trajectory has fallen into the low-energy basin at *A* ≈ 0.9–1.0, merely restoring load to its original value is insufficient; the separatrix continues to separate the collapsed state from the high-energy attractor.
2. **Rescue by pruning alone requires crossing the left fold**. Within this minimal model, re-entry into a monostable high-energy regime requires reducing effective load below the left fold (*A* ≲ 0.86). Without a quantitative mapping from synapse loss or axonal length reduction to *A*, it is uncertain whether physiologically plausible pruning magnitudes can achieve this threshold; current anatomical estimates suggest that it would require substantial loss of arbor.
3. **Hysteresis exists mathematically but may be limited physiologically**. The right fold is above most SNc neurons’ anatomical load, so the system is unlikely to traverse it in vivo. Instead, the practically relevant barrier is the left fold, which sets the minimum load required for bistability and thus for tipping-like behavior.

These considerations are consistent with the clinical observation that SNc degeneration, once initiated, proceeds effectively irreversibly, but they also highlight the need for quantitative calibration between anatomical pruning and the dimensionless load parameter *A*.

### S5. Summary of supplementary findings

The supplementary analyses demonstrate three key points:

1. **Not all biologically plausible architectures can generate bistability**. Specific nonlinear interactions—particularly nonlinear energy amplification—are required (21–23).
2. **The saddle-node bifurcation in the final minimal model is robust across one-at-a-time physiological parameter variations**, persisting under broad changes in *k*_*M*_, *L*_1_, and *C* (21–23). Robustness under correlated multi-parameter variation remains to be assessed, but the (*A, C*) stability map shows that bistability occupies a broad, continuous band rather than a fragile point-like region.
3. **Collapse is effectively irreversible in the physiological range of loads considered**, consistent with the clinical course and anatomical constraints in SNc neurons, provided that feasible pruning cannot reduce effective load below the left fold (1,3,15–17,24).

Together, these results strengthen the interpretation that selective vulnerability of SNc neurons can emerge from **the fundamental geometry of load-dependent energetic regulation**, without requiring uniquely fragile mitochondria or exclusive molecular insults (15–17,21–24).

### S8. Stability regimes in (*A, C*) space

**Supplementary Figure S8.**
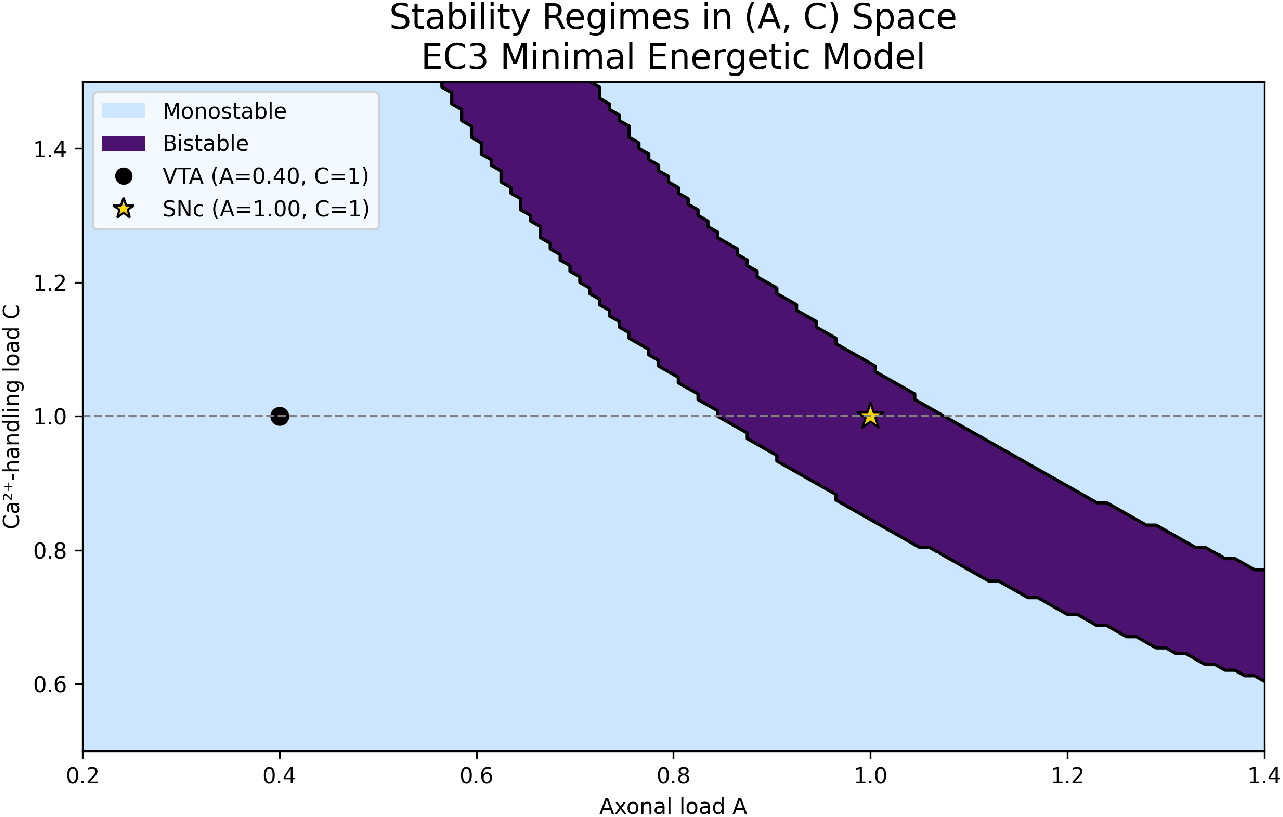
Stability regimes in (*A, C*) space for the EC3 minimal energetic model. Each point in the (*A, C*) plane was classified as monostable or bistable by numerically locating all equilibria in the physical domain [0, 1]^2^ and counting how many distinct fixed points exist (Methods, Section 7.8). Blue shading indicates **monostable** parameter combinations (a single equilibrium), and purple shading indicates **bistable** combinations (three equilibria: two stable fixed points and one saddle). The dashed horizontal line marks *C* = 1. The black circle marks the VTA-like condition (*A, C*) = (0.40, 1.0), which lies safely in the monostable region. The gold star marks the SNc-like condition (*A, C*) = (1.00, 1.0), which lies inside the bistable band. The bistable region forms an oblique band in (*A, C*) space: at **lower** Ca^2^ load *C*, higher axonal load *A* is required to induce bistability; at **higher** *C*, the same saddle-node structure appears at lower *A*. Thus, axonal arborization and Ca^2^-handling demand act as **multiplicative co-loads** that jointly determine whether a neuron resides in a monostable or bistable energetic regime.

### S9. Boundary vector field and positive invariance

**Supplementary Figure S9.**
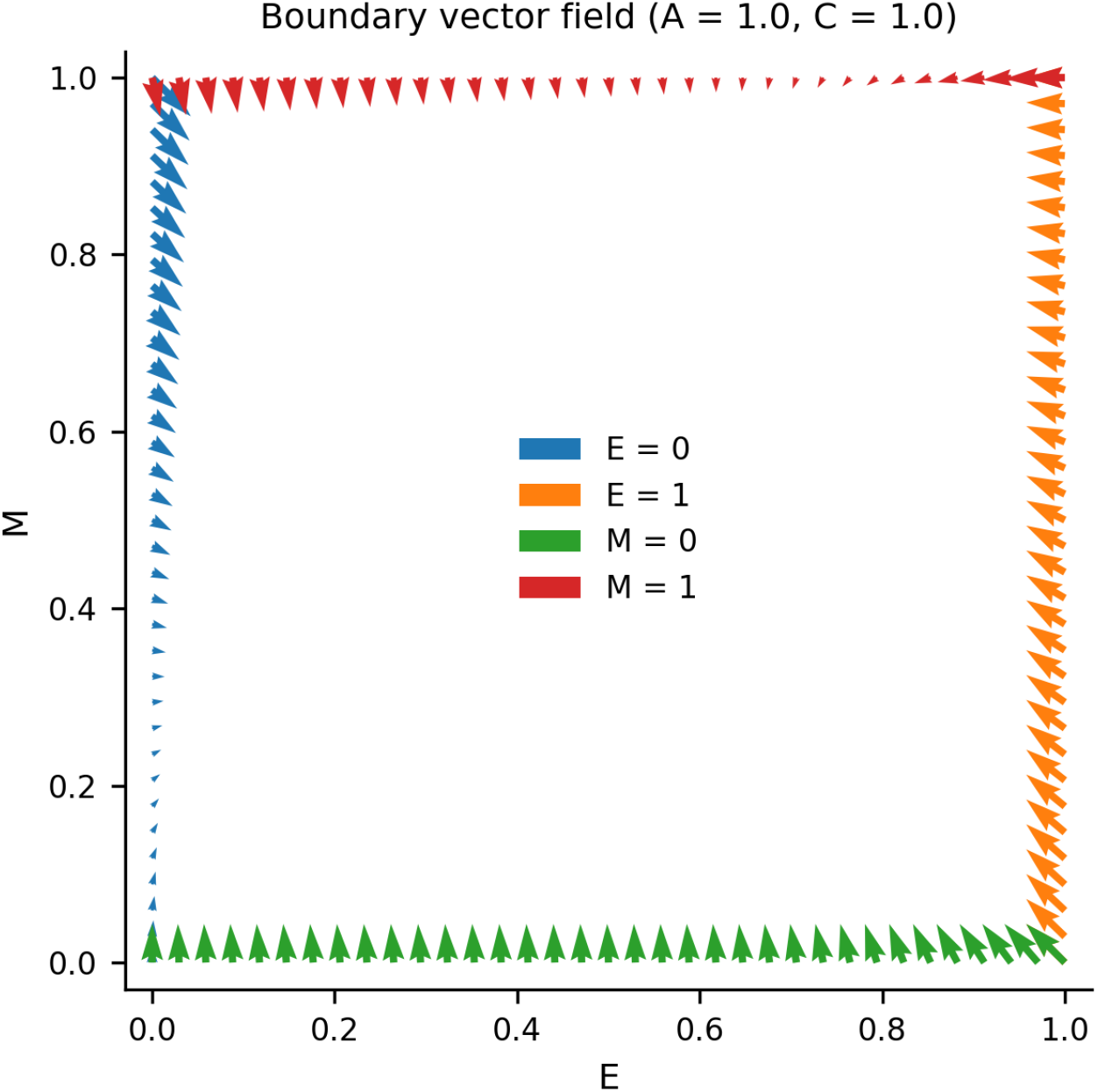
Vector field along the boundary of the physiological domain (*E, M*) ∈ [0, 1]^2^. Arrows show the direction of the ODE vector field along each edge of the square domain. At *E* = 0, vectors point inward with increasing *E*; at *E* = 1, they point inward with decreasing *E*; at *M* = 0, they point inward with increasing *M*; and at *M* = 1, they point inward with decreasing *M*. All boundary vectors point strictly into the interior (up to tangency at the corners), confirming that the physiological domain is positively invariant: trajectories that start in [0, 1]^2^ cannot leave it.

### S10. Joint (*E, M*) perturbation dynamics

**Supplementary Figure S10.**
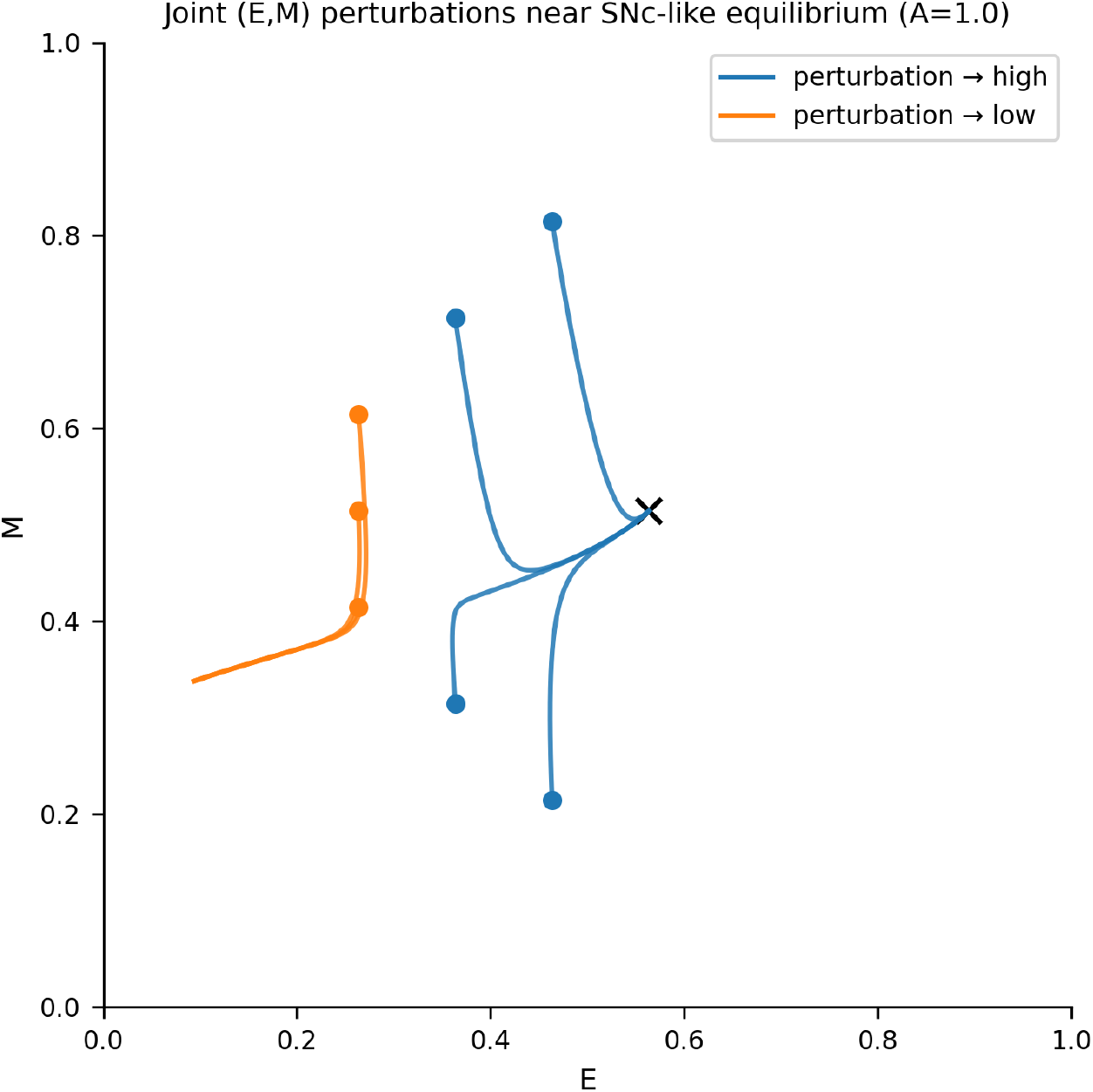
Trajectories following joint perturbations of the SNc-like equilibrium in (*E, M*) space. Starting from the high-energy SNc-like fixed point, we applied small perturbations in random directions within a ball of radius *δ* in (*E, M*) space and integrated the trajectories forward in time. Perturbations whose initial conditions lay above the separatrix (blue trajectories) returned to the high-energy attractor, whereas those below the separatrix (orange trajectories) converged to the low-energy attractor. The transition between outcomes occurs across a narrow geometric boundary: collapse is governed by the location of the separatrix, not by the specific direction of perturbation. Any multidimensional insult that pushes the system across this boundary produces irreversible energetic collapse in the model.

### S11. Bistability as a function of *A* and *k*_2_

**Supplementary Figure S11.**
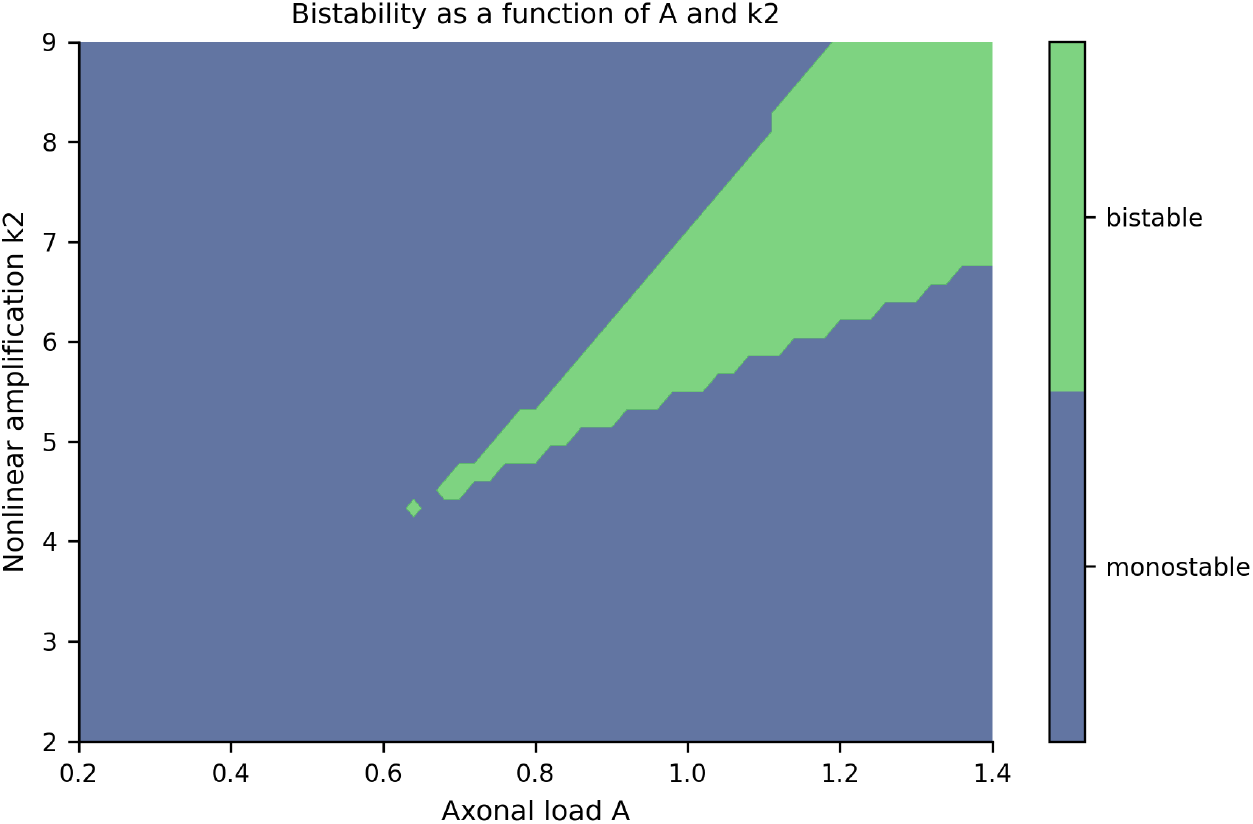
Bistable regimes in (*A, k*_2_) space. The axonal load *A* and nonlinear amplification parameter *k*_2_ were varied on a 61 × 51 grid, and each parameter pair was classified as monostable or bistable by counting equilibria in [0, 1]^2^. Blue shading indicates monostable parameter combinations; purple shading indicates bistable combinations. The bistable region persists across a broad range of *k*_2_ values (4–9) and maintains a smooth, diagonally oriented geometry. This demonstrates that bistability does not rely on fine-tuning the nonlinear amplification term, but is instead a structurally stable feature of the feedback motif.

### S12. Population heterogeneity in axonal load

**Supplementary Figure S12.**
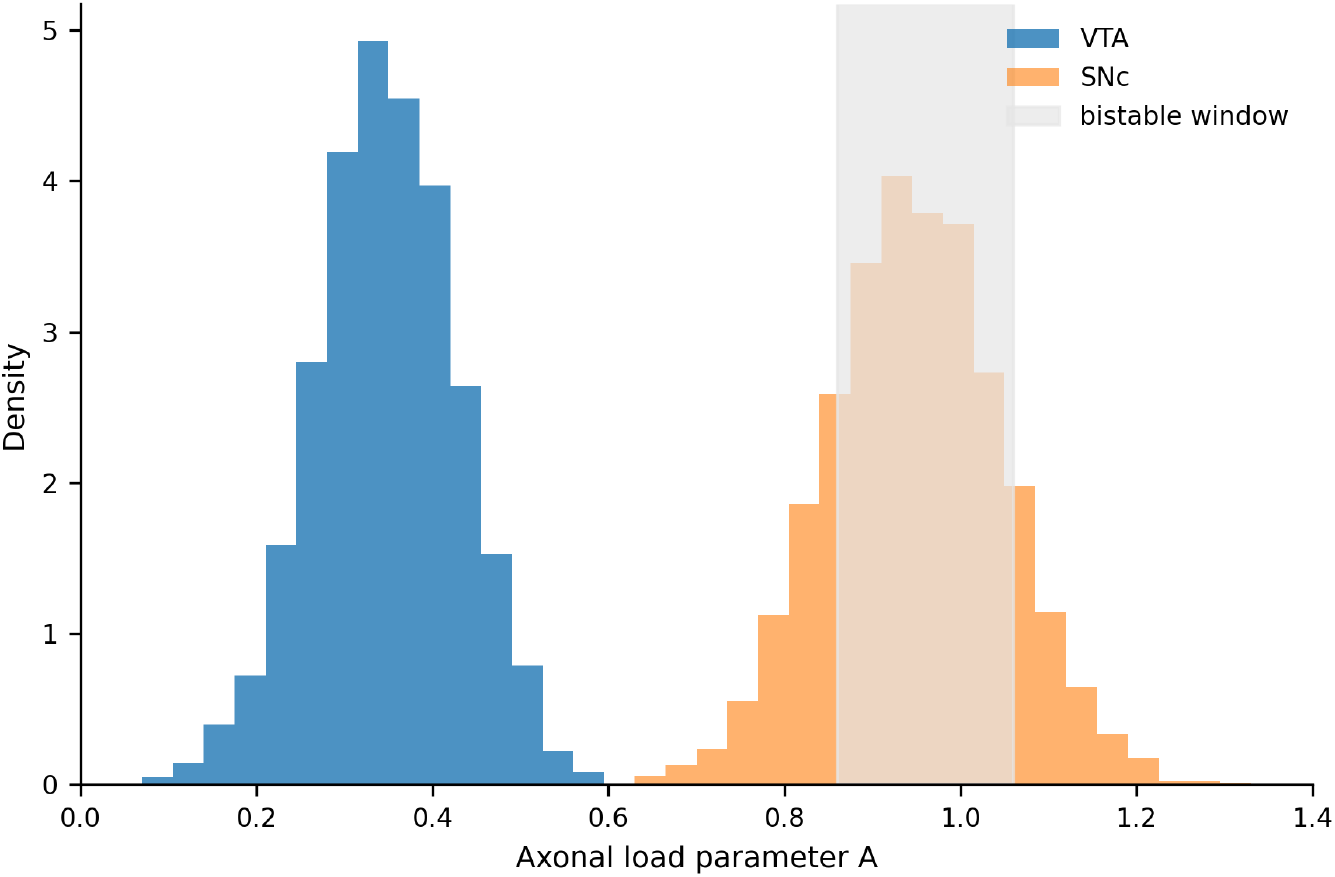
Synthetic population distributions of SNc and VTA neurons in axonal-load space. Histograms show the distributions of effective axonal load *A* for synthetic populations of 5,000 VTA neurons (blue) and 5,000 SNc neurons (gold), constructed using literature-reported ranges of arborization size and synapse number (1,3,15–17). The shaded vertical band marks the bistable window *A* ∈ [0.86, 1.06] from the main bifurcation analysis. VTA neurons cluster at low *A* (≈ 0.3–0.5), entirely outside the bistable window. SNc neurons cluster near *A* ≈ 0.9–1.0, with the majority falling inside the bistable window and a subset of high-arbor outliers approaching the right fold. Anatomical variability therefore preserves, rather than erases, the separation between stable (VTA) and vulnerable (SNc) regimes at the population level.

### S13. Solver robustness

**Supplementary Figure S13.**
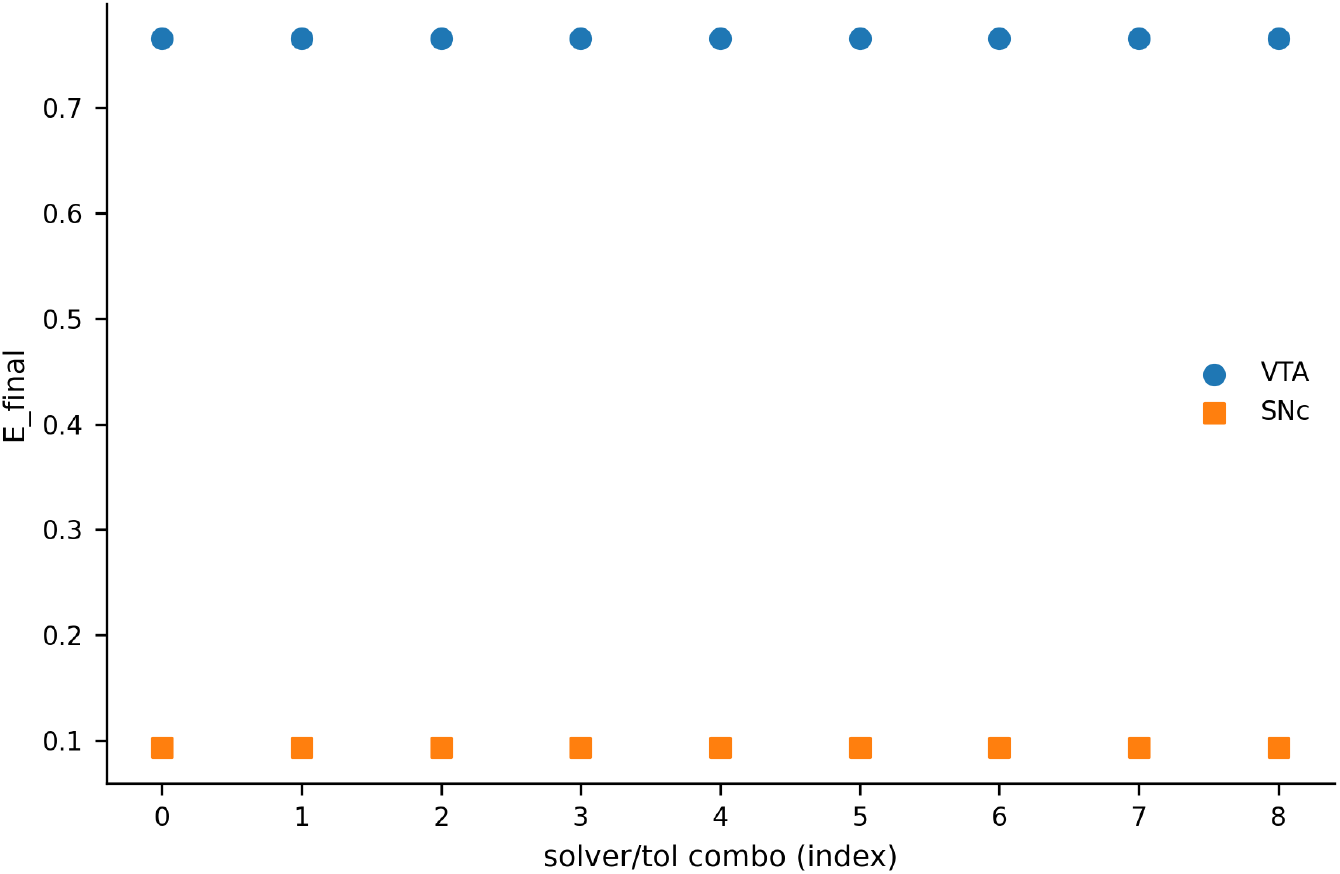
Robustness of regime classification across numerical solvers and tolerances. Final energy values *E*_final_ for VTA-like and SNc-like simulations are shown for nine solver–tolerance combinations: three integrators (RK45, RK23, BDF) crossed with three tolerance settings (default, 10× tighter, 10× looser). For each combination, trajectories were initialized near the high-energy equilibrium and subjected to the perturbation protocol described in Methods (Section 7.3). All combinations produced identical qualitative outcomes: VTA-like trajectories always returned to the high-energy attractor, and SNc-like trajectories either remained at or collapsed to the low-energy attractor depending on whether the perturbation crossed the separatrix. No solver produced spurious equilibria or oscillations. These results exclude the possibility that the bistable structure arises from numerical artifacts.

### Supplement S14. Collapse Is Reversible Only When Pruning Reduces Load Below the Left Fold

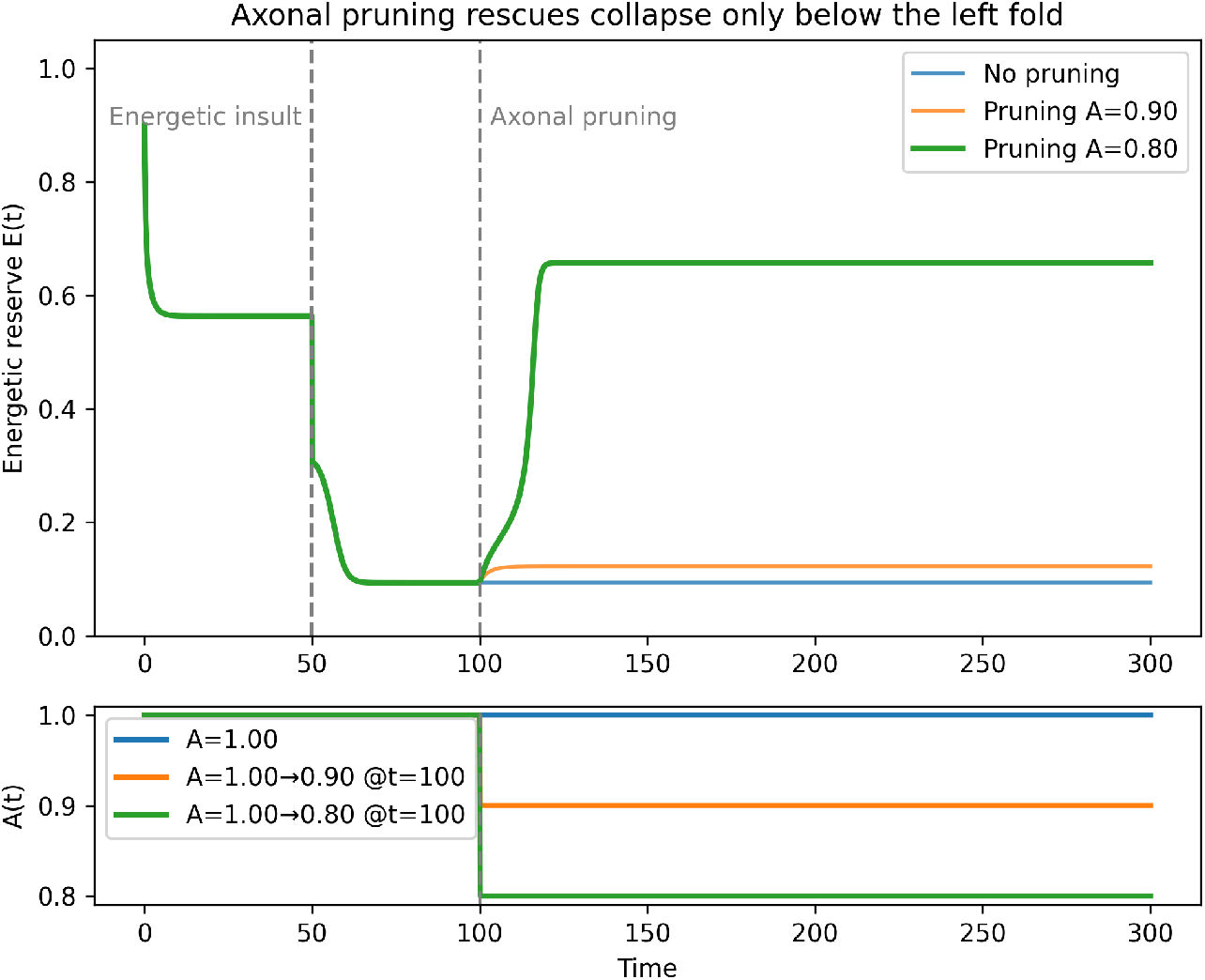

To test whether anatomical pruning can reverse an already-initiated energetic collapse, we simulated SNc-like neurons (baseline *A* = 1.00) subjected to the same brief energetic insult used in Figure 5 (resetting *E*(*t*) to 0.3 at *t* = 50). After the insult, axonal load was either left unchanged, reduced modestly to *A* = 0.90, or reduced substantially to *A* = 0.80. Because the bifurcation analysis identified a left saddle-node fold at *A* ≈ 0.86, these conditions place the system respectively **inside the bistable window, still inside the bistable window**, and **below the fold** where only a single high-energy equilibrium exists.

**Figure S14’s top panel** shows the resulting energetic trajectories.

- With **no pruning** or **moderate pruning to** *A* = 0.90, the system remains trapped in the collapsed low-energy basin. In both cases the low-energy attractor persists, and the trajectory cannot cross the separatrix back into the high-energy basin.
- In contrast, **pruning to** *A* = 0.80 eliminates the collapsed fixed point entirely, forcing the trajectory into the unique high-energy equilibrium. This behavior is independent of initial post-insult mitochondrial state because the monostable regime admits no collapsed attractor.

These results confirm the **geometric prediction** from the bifurcation analysis: **Once collapse occurs, pruning can rescue the neuron only if effective load is reduced below the left saddle-node**. Pruning inside the bistable range (*A* ∈ [0.86, 1.06]) cannot reverse collapse because the low-energy attractor continues to exist and remains dynamically accessible.

**Figure S14’s bottom panel** displays the piecewise-constant axonal load schedules *A*(*t*) used in the simulations. In all cases pruning occurred at *t* = 100, after the insult-induced trajectory had already entered the neighborhood of the collapsed state. The contrast between the *A* = 0.90 and *A* = 0.80 trajectories demonstrates that **the success or failure of pruning is determined not by timing or insult magnitude but by whether the parameter shift exits the bistable band in** (*A, C*)**-space** (cf. Supplementary Figure S8).

Together, these findings highlight a practical implication of the model: **reversal of collapse requires leaving the bistable regime entirely**. Within the anatomical ranges reported for SNc neurons, this would require unusually large reductions in arborization or concurrent modifications of additional parameters (e.g., *k*_*M*_, *C*) capable of shifting the fold. As a result, the model predicts that typical levels of axonal pruning alone are insufficient to restore collapsed SNc neurons, consistent with the clinical irreversibility of dopaminergic degeneration.

### Supplement S15. Stochastic Separatrix Crossing Distinguishes Fluctuation-Driven From Insult-Driven Collapse

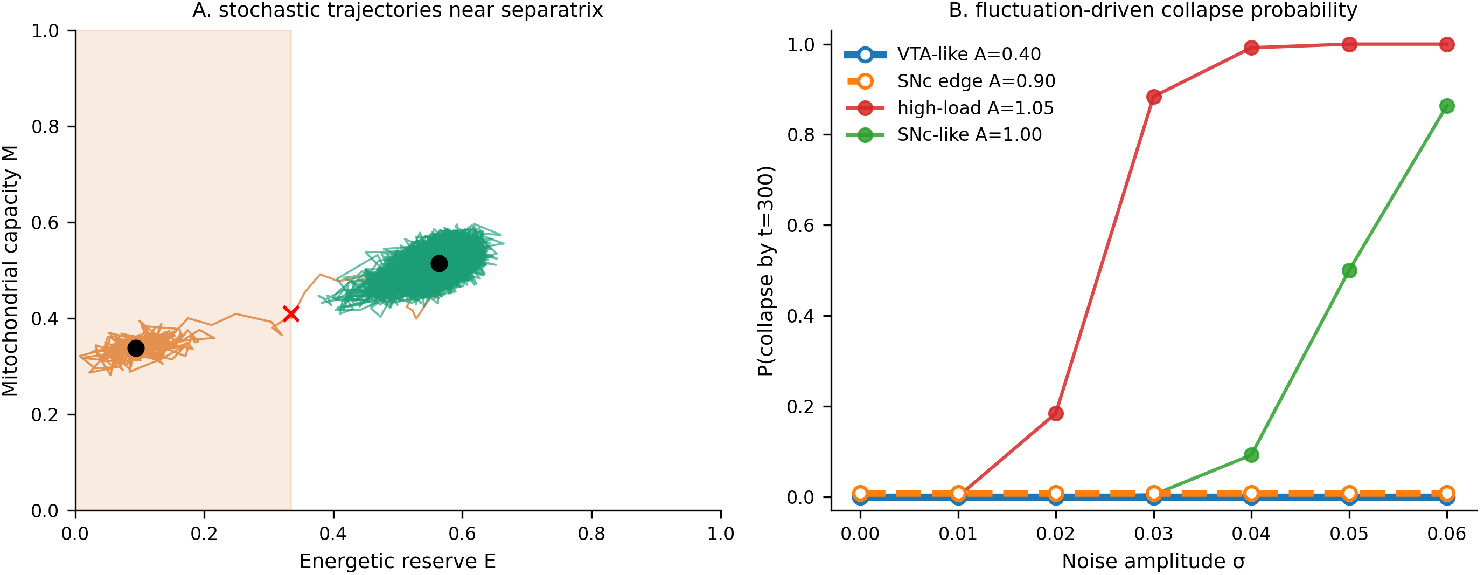

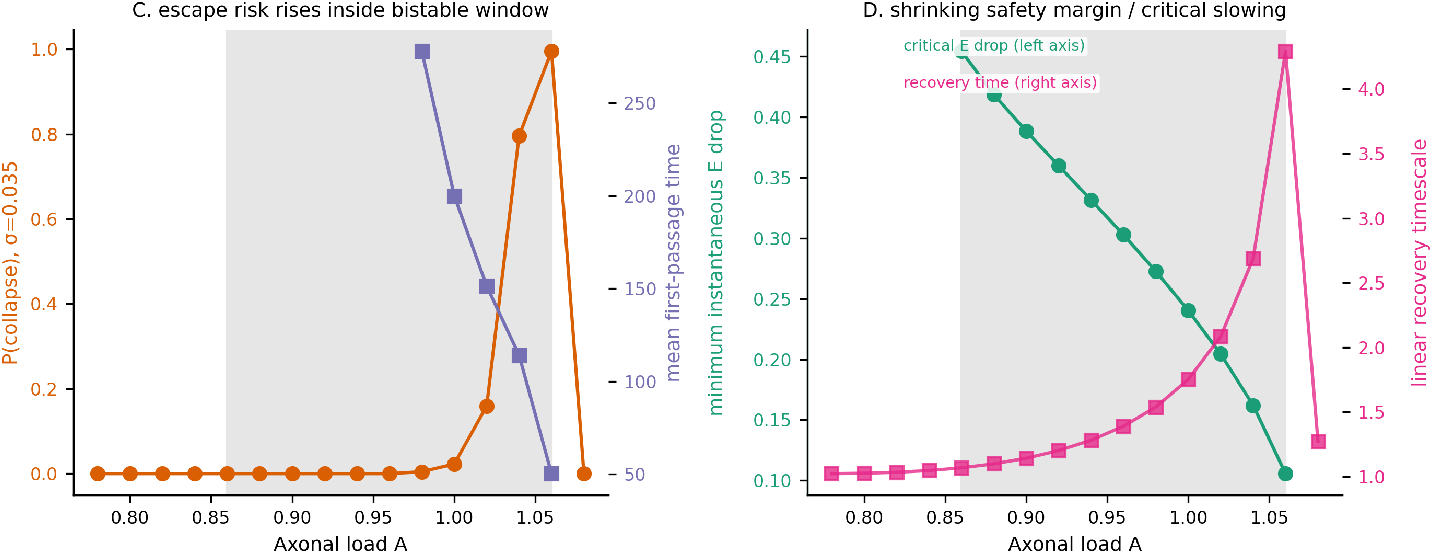

To test whether the EC3 separatrix can be crossed by intrinsic fluctuations rather than by an imposed deterministic insult, we simulated a stochastic version of the model with additive noise in both *E* and *M* (Methods, Section 7.12). Each trajectory was initialized at the high-energy equilibrium for its axonal load *A*, and collapse was defined as persistent crossing to the low-energy side of the saddle-coordinate proxy. Because no irreversible collapsed attractor exists outside the bistable regime, VTA-like monostable conditions were assigned zero irreversible escape probability.

**Figure S15A** shows representative noisy trajectories at *A* = 1.00. Most trajectories fluctuate near the high-energy attractor, but sufficiently large excursions can cross the saddle-side boundary and relax toward the low-energy attractor. **Figure S15B** sweeps the dimensionless noise amplitude *σ*: VTA-like loads remain protected across the tested range, whereas SNc-like and high-load SNc conditions show sharply increasing collapse probability as the high-energy basin narrows. At *A* = 1.05, near the right fold, even modest noise produces frequent first-passage events.

**Figure S15C** holds *σ* = 0.035 fixed and varies *A*. Collapse probability rises steeply near the high-load edge of the bistable window, while mean first-passage time falls. **Figure S15D** provides the deterministic counterpart: the minimum instantaneous energy drop required to enter the collapsed basin decreases as *A* increases, and the linear recovery timescale grows near the fold. Together these results distinguish two mechanisms within the same geometry. Far from the separatrix, collapse requires a discrete insult large enough to cross the basin boundary; near the fold, intrinsic fluctuations alone can produce separatrix crossing on finite timescales.

The stochastic analysis is not a calibrated disease-progression model, because *σ* is dimensionless and not fitted to measured ATP or mitochondrial fluctuations. Its value is therefore comparative: it shows how the EC3 geometry predicts a graded transition from insult-dominated collapse at lower SNc-like loads to fluctuation-sensitive collapse near the fold, while preserving VTA-like protection in the monostable regime.

### Supplement S17. Model-Form, Population, and Insult-Normalization Sensitivity

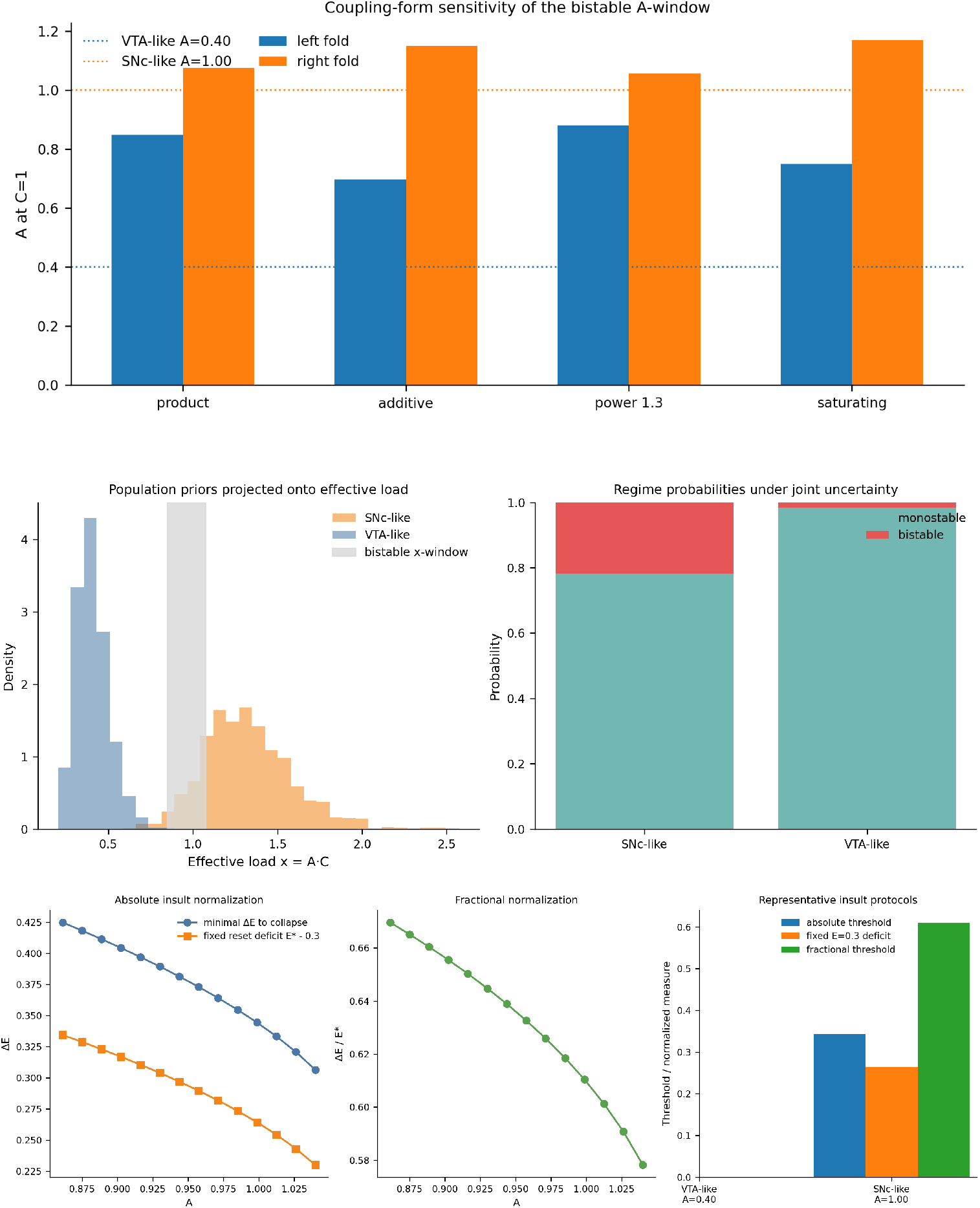

To probe which claims depend on the exact algebraic form of the model and which remain robust under reasonable no-new-data perturbations, we ran three sensitivity checks: alternative A–C coupling forms, synthetic VTA/SNc uncertainty propagation, and insult-normalization comparisons. All preserve the qualitative bistable geometry. The mapped A-window shifts with the coupling choice, VTA-like samples remain mostly monostable, SNc-like samples retain a substantial bistable fraction, and fixed absolute *E* resets are not equivalent to fractional thresholds across populations.

### Supplement S18. Parameter Sensitivity of the Bistable Fold Window

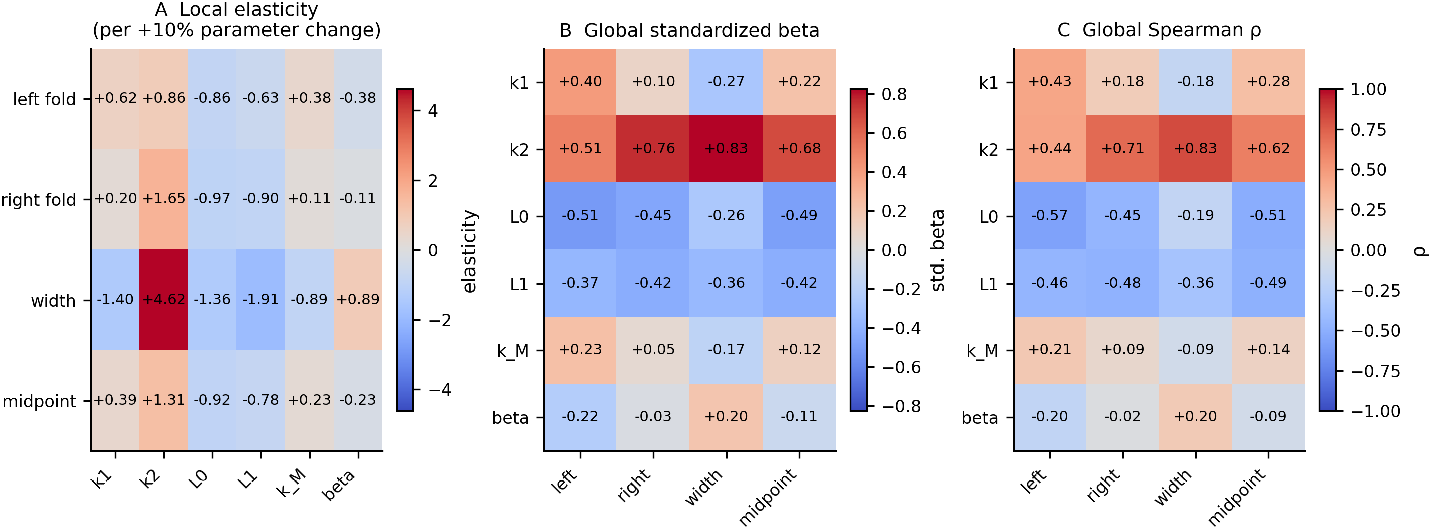

A local one-at-a-time scan and a global Monte Carlo ranking show that the baseline fold window spans *A* ≈ 0.849 to 1.075 (width ≈ 0.226, midpoint ≈ 0.962). The width is most sensitive to *k*_2_, with secondary contributions from *L*_1_, *L*_0_, and *k*_1_; *k*_*M*_ and *β* are weaker modulators. The global ranking matches the local picture and explains most of the variance in fold location and width (all *R*^2^ ≳ 0.94).

### Supplement S19. Mechanism Summary of Fold Geometry and Sensitivity

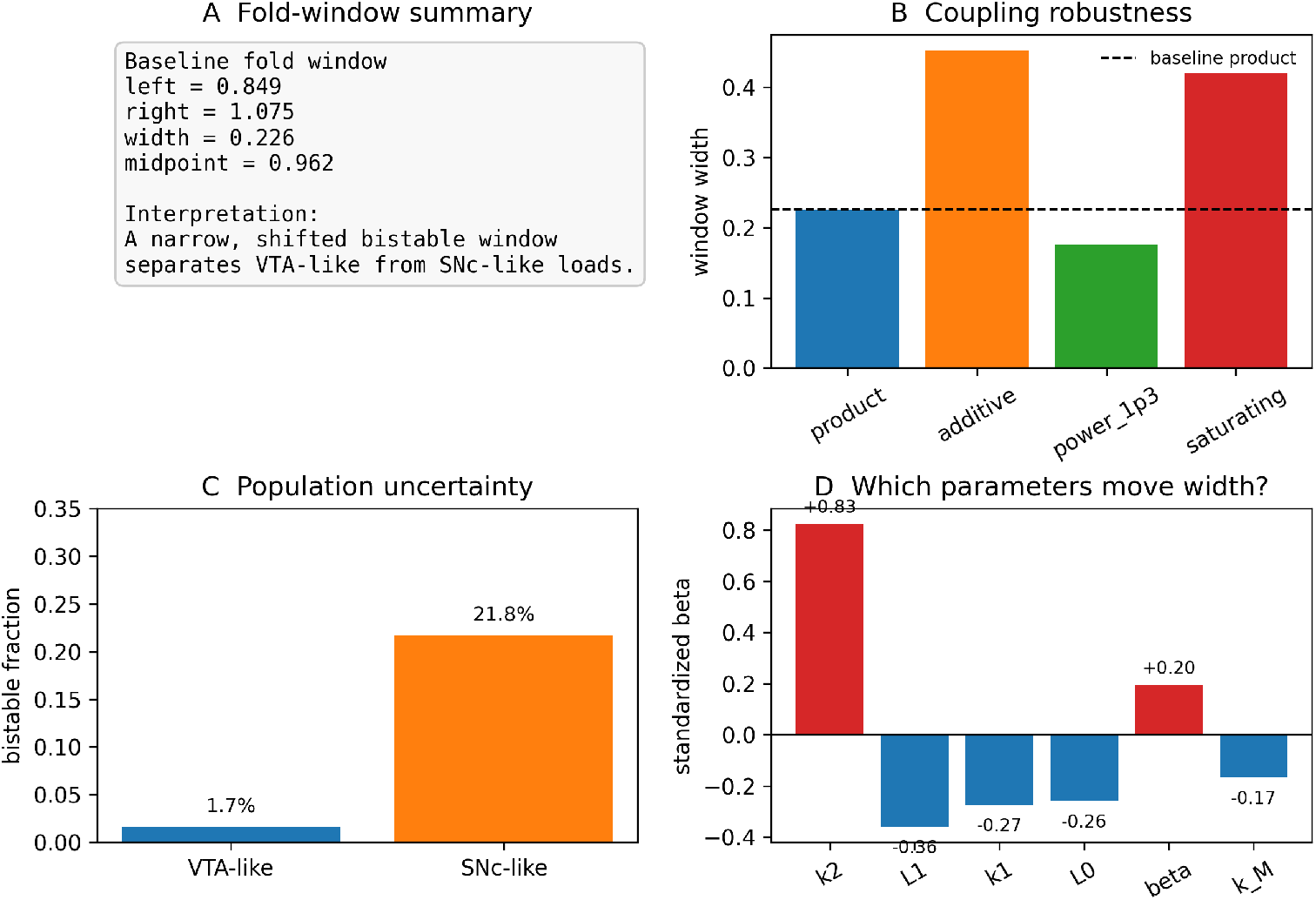

To make the S17–S18 results easier to read at a glance, we assembled a compact mechanism summary. The summary collects the baseline fold window, the coupling-form window widths, the VTA/SNc bistability fractions under synthetic uncertainty, and the dominant width-sensitivity parameters in one view. The synthesis does not add new biological assumptions; instead, it emphasizes the main operational takeaway: the bistable window is narrow but not fragile, VTA-like loads remain near the monostable regime, SNc-like loads remain inside the collapse-prone regime, and the width of the tipping window is dominated by the nonlinear amplification term *k*_2_ with secondary contributions from *L*_1_ and *L*_0_.

### Supplement S20. Minimal Two-Compartment Extension of the EC3 Model

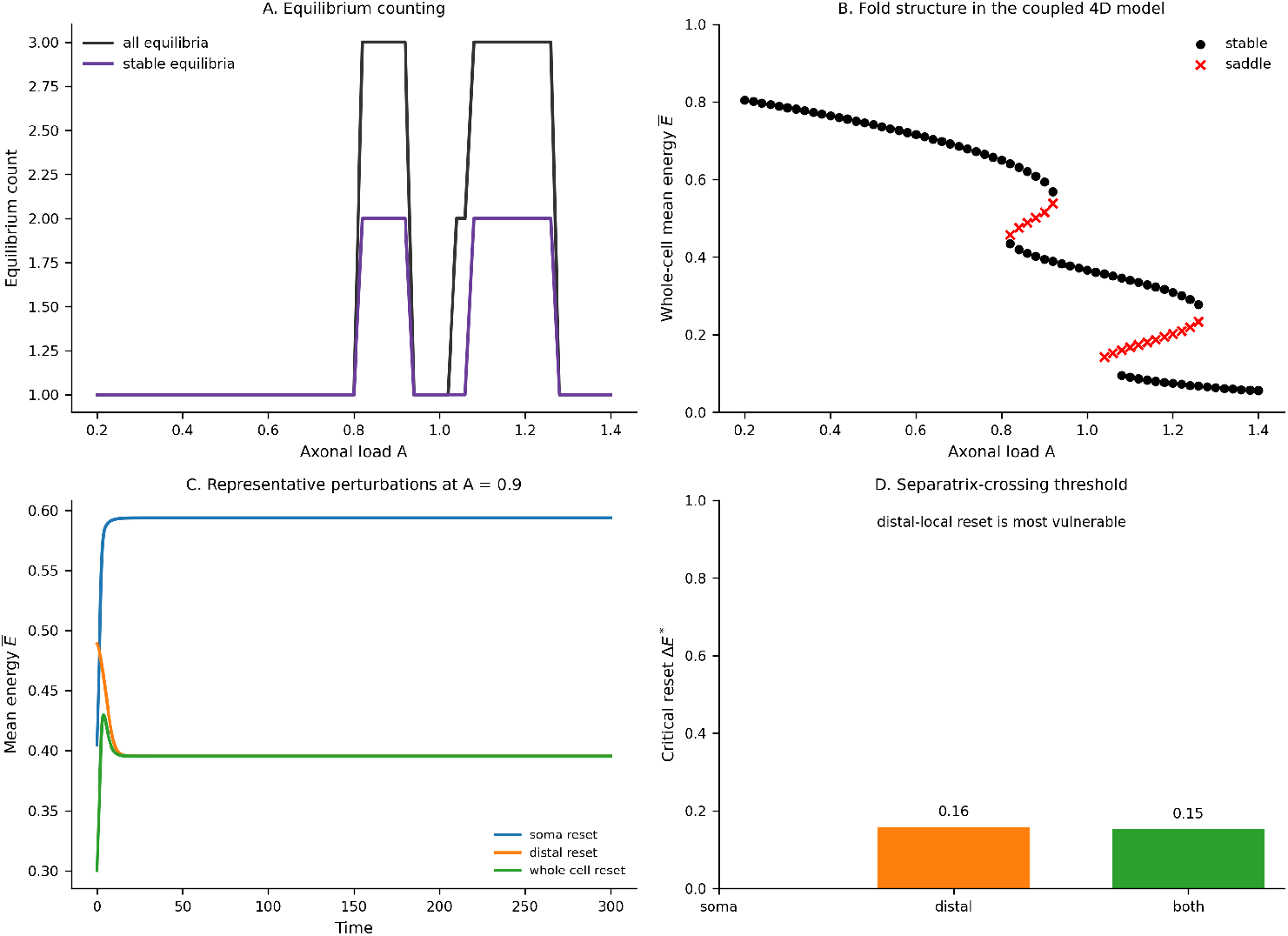

To test whether the tipping geometry survives a minimal spatial compartmentalization of the arbor, we split the neuron into soma and distal-arbor energy/mitochondrial compartments with exchange terms and placed most of the load on the distal compartment. The coupled 4D system still supports bistability, but the window is reshaped into two lobes rather than a single uninterrupted band: a lower-load lobe around *A* ≈ 0.82–0.92 and a higher-load lobe around *A* ≈ 1.08–1.26, separated by a monostable gap near *A* ≈ 1.0–1.06. Representative perturbations show that soma-local resets can recover while distal-local or whole-cell resets can cross the separatrix and collapse. This indicates that the load-induced tipping geometry is not an artifact of whole-cell homogenization, although the exact fold structure becomes compartment-sensitive.

### Supplement S21. Blind Prior-Predictive Calibration Stress Test

The blind prior-predictive / leave-one-population-out calibration reduced but did not eliminate the circularity concern. Under the reference blind prior, VTA-like samples remained monostable with very high probability, but SNc-like samples remained bistable only in a minority of draws, and the leave-one-population-out holdout remained prior-sensitive. We therefore treat this check as a blocker note rather than a standalone claim-strengthening result, and we do not use it to overstate de-circularization of the SNc/VTA mapping.

## References

1. Matsuda W, Furuta T, Nakamura KC, Hioki H, Fujiyama F, Arai R, Kaneko T. Single nigrostriatal dopaminergic neurons form widely spread and highly dense axonal arborizations in the neostriatum. J Neurosci. 2009;29(2):444–453. doi:10.1523/JNEUROSCI.4029-08.2009

2. Bolam JP, Hanley JJ, Booth PA, Bevan MD. Synaptic organisation of the basal ganglia. J Anat. 2000;196(4):527–542. doi:10.1046/j.1469-7580.2000.19640527.x

3. Gauthier J, Parent M, Lévesque M, Parent A. The axonal arborization of single nigrostriatal neurons in rats. Brain Res. 1999;834(1–2):228–232. doi:10.1016/S0006-8993(99)01573-5

4. Pan WX, Mao T, Dudman JT. Inputs to the dorsal striatum of the mouse reflect the parallel circuit architecture of the forebrain. Front Neuroanat. 2010;4:147. doi:10.3389/fnana.2010.00147

5. Surmeier DJ, Guzman JN, Sanchez-Padilla J, Goldberg JA. What causes the death of dopaminergic neurons in Parkinson’s disease? Prog Brain Res. 2010;183:59–77. doi:10.1016/S0079-6123(10)83004-3

6. Guzman JN, Sánchez-Padilla J, Chan CS, Surmeier DJ. Robust pacemaking in substantia nigra dopaminergic neurons. J Neurosci. 2009;29(35):11011–11019. doi:10.1523/JNEUROSCI.2519-09.2009

7. Chan CS, Guzman JN, Ilijic E, et al. ‘Rejuvenation’ protects neurons in mouse models of Parkinson’s disease. Nature. 2007;447(7148):1081–1086. doi:10.1038/nature05865

8. Zampese E, Wokosin DL, Gonzalez-Rodriguez P, Guzman JN, et al. Ca^2^ channels couple spiking to mitochondrial metabolism in substantia nigra dopaminergic neurons. Sci Adv. 2022;8(37):eabp8701. doi:10.1126/sciadv.abp8701

9. Surmeier DJ. Calcium, ageing, and neuronal vulnerability in Parkinson’s disease. Lancet Neurol. 2007;6(10):933–938. doi:10.1016/S1474-4422(07)70246-6

10. Exner N, Lutz AK, Haass C, Winklhofer KF. Mitochondrial dysfunction in Parkinson’s disease: molecular mechanisms and pathophysiological consequences. EMBO J. 2012;31(14):3038–3062. doi:10.1038/emboj.2012.170

11. Grünewald A, Rygiel KA, Zsurka G, et al. Mitochondrial DNA depletion in substantia nigra neurons in Parkinson disease. Ann Neurol. 2016;79(3):366–378. doi:10.1002/ana.24571

12. Burbulla LF, Song P, Mazzulli JR, et al. Dopamine oxidation mediates mitochondrial and lysosomal dysfunction in Parkinson’s disease. Science. 2017;357(6357):1255–1261. doi:10.1126/science.aam9080

13. Schapira AHV, Cooper JM, Dexter D, Clark JB, Jenner P, Marsden CD. Mitochondrial complex I deficiency in Parkinson’s disease. J Neurochem. 1990;54(3):823–827. doi:10.1111/j.1471-4159.1990.tb02325.x

14. Bose A, Beal MF. Mitochondrial dysfunction in Parkinson’s disease. J Neurochem. 2016;139(Suppl 1):216–231. doi:10.1111/jnc.13731

15. Pacelli C, Giguère N, Bourque M-J, Lévesque M, Slack RS, Trudeau L-É. Elevated mitochondrial bioenergetics and axonal arborization size are key contributors to the vulnerability of dopamine neurons. Curr Biol. 2015;25(18):2349–2360. doi:10.1016/j.cub.2015.07.050

16. Giguère N, Burke Nanni S, Trudeau L-E. On cell loss and selective vulnerability of neuronal populations in Parkinson’s disease. Front Neurol. 2018;9:455. doi:10.3389/fneur.2018.00455

17. Surmeier DJ. Determinants of dopaminergic neuron loss in Parkinson’s disease. FEBS J. 2018;285(19):3657–3668. doi:10.1111/febs.14607

18. Wong YC, Krainc D. -Synuclein toxicity in neurodegeneration: mechanism and therapeutic strategies. Nat Med. 2017;23(2):1–13. doi:10.1038/nm.4269

19. Conway KA, Harper JD, Lansbury PT. Accelerated in vitro fibril formation by a mutant-synuclein linked to early-onset Parkinson disease. Nat Med. 1998;4(11):1318–1320. doi:10.1038/3311

20. Cookson MR. The role of leucine-rich repeat kinase 2 (LRRK2) in Parkinson’s disease. Nat Rev Neurosci. 2010;11(12):791–797. doi:10.1038/nrn2935

21. Strogatz SH. Nonlinear Dynamics and Chaos. 2nd ed. CRC Press; 2018. doi:10.1201/9780429492563

22. Scheffer M, Carpenter SR, Lenton TM, et al. Anticipating critical transitions. Science. 2012;338(6105):344–348. doi:10.1126/science.1225244

23. Kuehn C. A mathematical framework for critical transitions: normal forms, variance and applications. J Nonlinear Sci. 2011;23(3):457–510. doi:10.1007/s00332-012-9158-x

24. Harris JJ, Jolivet R, Attwell D. Synaptic energy use and supply. Neuron. 2012;75(5):762–777. doi:10.1016/j.neuron.2012.08.019

25. Attwell D, Laughlin SB. An energy budget for signaling in the grey matter of the brain. J Cereb Blood Flow Metab. 2001;21(10):1133–1145. doi:10.1097/00004647-200110000-00001

26. Virtanen P, Gommers R, Oliphant TE, et al. SciPy 1.0: fundamental algorithms for scientific computing in Python. Nat Methods. 2020;17(3):261–272. doi:10.1038/s41592-019-0686-2

27. Hunter JD. Matplotlib: A 2D graphics environment. Comput Sci Eng. 2007;9(3):90–95. doi:10.1109/MCSE.2007.55

28. Harris CR, Millman KJ, van der Walt SJ, et al. Array programming with NumPy. Nature. 2020;585(7825):357–362. doi:10.1038/s41586-020-2649-2

29. Prensa L, Parent A. The nigrostriatal pathway in the rat: a single-axon study. J Neurosci. 2001;21(18):7247–7260. doi:10.1523/JNEUROSCI.21-18-07247.2001

30. Bolam JP, Pissadaki EK. Living on the edge with too many mouths to feed: why dopamine neurons die. Mov Disord. 2012;27(12):1478–1483. doi:10.1002/mds.25135

31. Pissadaki EK, Bolam JP. The energy cost of action potential propagation in dopamine neurons: clues to susceptibility in Parkinson’s disease. Front Comput Neurosci. 2013;7:13. doi:10.3389/fncom.2013.00013

32. Muddapu VR, Chakravarthy VS, Ravindran B, Moustafa AA. Influence of energy deficiency on the subcellular processes of substantia nigra pars compacta cell for understanding Parkinsonian neurodegeneration. Sci Rep. 2021;11:1754. doi:10.1038/s41598-021-81185-9

33. Guzman JN, Sanchez-Padilla J, Wokosin D, Kondapalli J, Ilijic E, Schumacker PT, Surmeier DJ. Oxidant stress evoked by pacemaking in dopaminergic neurons is attenuated by DJ-1. Nature. 2010;468(7324):696–700. doi:10.1038/nature09536

34. Amini B, Clark JW Jr, Canavier CC. Calcium dynamics underlying pacemaker-like and burst firing oscillations in midbrain dopaminergic neurons: a computational study. J Neurophysiol. 1999;82(5):2249–2261. doi:10.1152/jn.1999.82.5.2249

35. Wilson CJ, Callaway JC. Coupled oscillator model of the dopaminergic neuron of the substantia nigra. J Neurophysiol. 2000;83(5):3084–3100. doi:10.1152/jn.2000.83.5.3084

36. Kuznetsova AY, Huertas MA, Kuznetsov AS, Paladini CA, Canavier CC. Regulation of firing frequency in a computational model of a midbrain dopaminergic neuron. J Comput Neurosci. 2010;28(3):389–403. doi:10.1007/s10827-010-0222-y

37. Philippart F, Destreel G, Merino-Sepúlveda P, Henny P, Engel D, Seutin V. Differential somatic Ca2+ channel profile in midbrain dopaminergic neurons. J Neurosci. 2016;36(27):7234–7245. doi:10.1523/JNEUROSCI.0459-16.2016

38. Cortassa S, Aon MA, Marbán E, Winslow RL, O’Rourke B. An integrated model of cardiac mitochondrial energy metabolism and calcium dynamics. Biophys J. 2003;84(4):2734–2755. doi:10.1016/S0006-3495(03)75079-6

39. Magnus G, Keizer J. Minimal model of beta-cell mitochondrial Ca2+ handling. Am J Physiol. 1997;273(2):C717–C733. doi:10.1152/ajpcell.1997.273.2.C717

40. Magnus G, Keizer J. Model of beta-cell mitochondrial calcium handling and electrical activity. II. Mitochondrial variables. Am J Physiol. 1998;274(4):C1174–C1184. doi:10.1152/ajpcell.1998.274.4.C1174

41. Bertram R, Gram Pedersen M, Luciani DS, Sherman A. A simplified model for mitochondrial ATP production. J Theor Biol. 2006;243(4):575–586. doi:10.1016/j.jtbi.2006.07.019

42. Ge P, Dawson VL, Dawson TM. PINK1 and Parkin mitochondrial quality control: a source of regional vulnerability in Parkinson’s disease. Mol Neurodegener. 2020;15:20. doi:10.1186/s13024-020-00367-7

43. Smith H. Monotone Dynamical Systems: An Introduction to the Theory of Competitive and Cooperative Systems. Providence, RI: American Mathematical Society; 2008. doi:10.1090/surv/041

44. Kuznetsov YA. Elements of Applied Bifurcation Theory. New York: Springer; 2004. doi:10.1007/978-1-4757-3978-7

